# Target Identification Under High Levels of Amplitude, Size, Orientation and Background Uncertainty

**DOI:** 10.1101/2024.08.30.610264

**Authors:** Can Oluk, Wilson S. Geisler

**Affiliations:** Center for Perceptual Systems and Department of Psychology, University of Texas at Austin; Laboratory of Psychophysics, Brain Mind Institute, École Polytechnique Fédérale de Lausanne (EPFL), Lausanne, Switzerland

## Abstract

Many natural tasks require the visual system to classify image patches accurately into target categories, including the category of no target. Natural target categories often involve high levels of within-category variability (uncertainty), making it challenging to uncover the underlying computational mechanisms. Here, we describe these tasks as identification from a set of exhaustive, mutually exclusive target categories, each partitioned into mutually exclusive subcategories. We derive the optimal decision rule and present a computational method to simulate performance for moderately large and complex tasks. We focus on the detection of an additive wavelet target in white noise with five dimensions of stimulus uncertainty: target amplitude, orientation, scale, background contrast, and spatial pattern. We compare the performance of the ideal observer with various heuristic observers. We find that a properly normalized heuristic MAX observer (SNN-MAX) approximates optimal performance. We also find that a convolutional neural network trained on this task approaches but does not reach optimal performance, even with extensive training.

We measured human performance on a task with three of these dimensions of uncertainty (orientation, scale, and background pattern). Results show that the pattern of hits and correct rejections for the ideal and SNN-MAX observers (but not a simple MAX observer) aligns with the data. Additionally, we measured performance under low uncertainty (without scale and orientation uncertainty) and found that the effect of uncertainty on the performance is smaller than any of the models predicted. This smaller-than-expected effect can largely be explained by including biologically plausible levels of intrinsic position uncertainty.

**Precis:** We describe target identification tasks in terms of mutually exclusive categories and subcategories and derive the optimal decision rule. Simulations of ideal and heuristic observers were compared to human data under high and low levels of extrinsic uncertainty.

## Introduction

In many natural tasks, the goal is to correctly identify an image as belonging to one out of some number of specific categories. For example, a natural task might be to indicate whether or not an image contains an apple (Figure 1a). Natural categories are generally quite complex because many different images fall into the same category; the images in Figure 1a are only a small subset of the images that would fall into the category of “target present.” Such high levels of within category variability (uncertainty) make natural categorization tasks difficult to perform and make it difficult to understand how and how well the human visual system performs the tasks.

**Figure 1.**
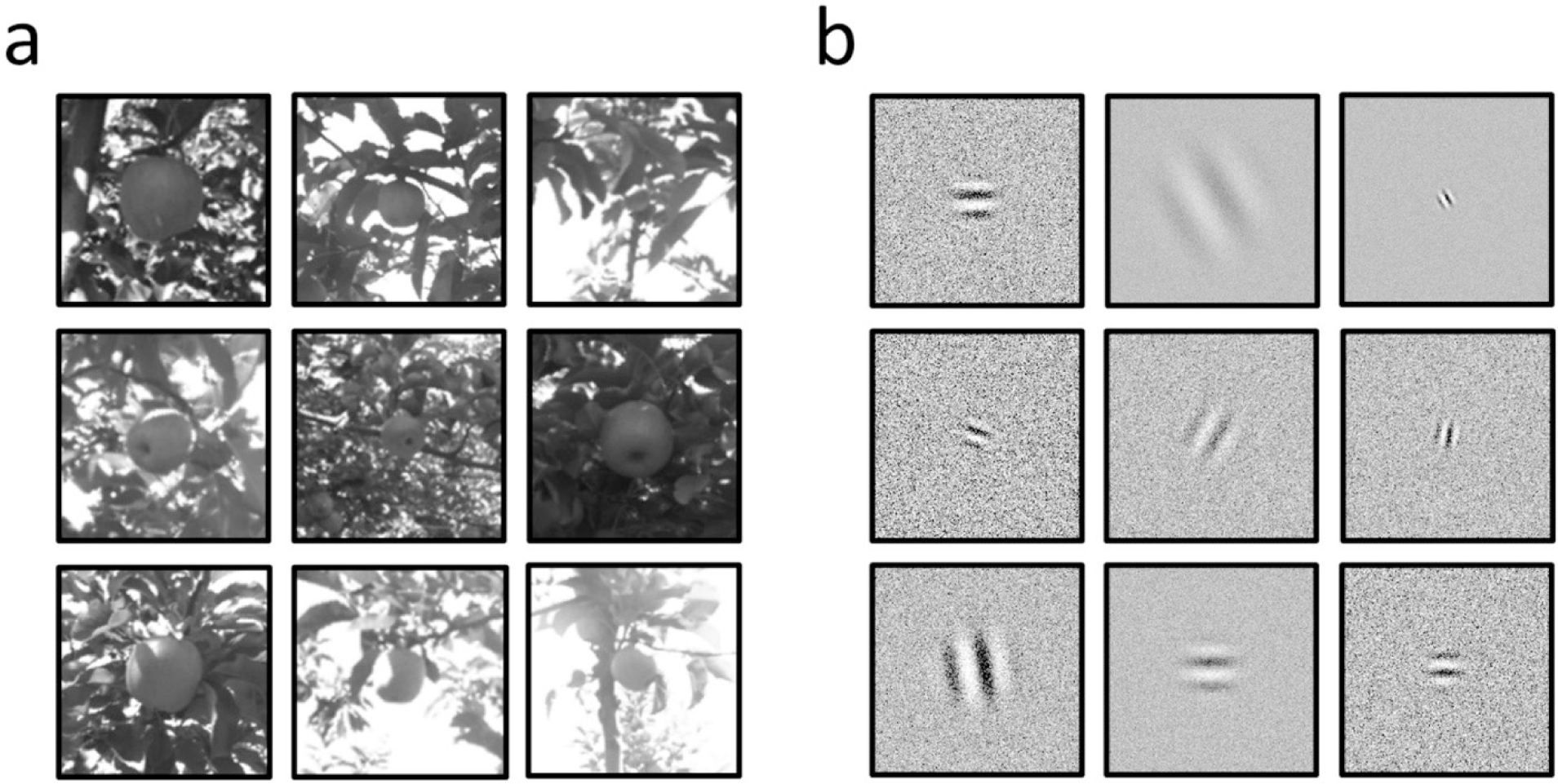
Identification under high levels of extrinsic uncertainty **a**. Multiple images of an apple (at the center) are shown under naturalistic settings. The variations in the properties of the apple (e.g., orientation, scale, contrast) and the background it appears against (e.g., contrast, configuration, brightness) demonstrate the vastness of the variation the visual system faces for the detection of an apple (Bargoti & Underwood,2017). **b**. Multiple images of a wavelet target in a white noise background. These mimic some of the natural extrinsic uncertainty by randomly picking target amplitude, orientation, scale, and background contrast from a broad range, while using a different sample of white noise background for each image, which also causes the background pattern (configuration) to randomly vary.

To gain some understanding of how and how well humans perform natural categorization tasks it is possible to use laboratory tasks that contain some of the variability that occurs in natural tasks (Davis et al., 1983; Eckstein & Abbey, 2001; Han & Baek, 2020; for reviews: Cohn & Lasley, 1986; Eckstein, 2011). For example, Figure 1b shows a small subset of the stimuli in a task where the goal is to indicate whether or not the image contains a wavelet target. The target-present images vary widely because of random variation in the amplitude, scale, and orientation of the wavelet and because of random variation in the spatial pattern and contrast of the noise background.

An advantage of such laboratory tasks is that the stimuli and prior probabilities are known precisely and hence it is possible, in principle, to determine the computations and performance of the ideal Bayesian observer in the categorization task. Such ideal observers provide insight into the computational principles needed for specific tasks and can serve as a benchmark for evaluating the performance of human and artificial visual systems (Green & Swets, 1966; Geisler, 2003; Geisler, 2011; Burge, 2020; Reith & Wandell, 2020).

The simplest and most direct approach for deriving and evaluating these Bayesian ideal observers is to partition the categories into mutually exclusive subcategories whose likelihoods can be directly calculated. These likelihoods can then be combined to determine the ideal observer’s response to a stimulus. A weakness of this direct approach is that for some tasks the number of mutually exclusive subcategories becomes too large for simulation of the ideal observer’s responses, given typical computational resources.

Here we first describe this direct simulation approach for arbitrary numbers of categories and subcategories. We then describe a computational method that allows simulation of performance for relatively large and complex tasks. We illustrate this approach for a binary categorization task with five dimensions of stimulus uncertainty (target amplitude, target orientation, target scale, background contrast, background spatial pattern; see Figure 1b) and compare the performance of the ideal observer for this task with that of heuristic observers including ones that utilize max operation, and a convolutional neural network (CNN) trained on this task. Finally, we describe measurements of human performance on a task with three of these dimensions of stimulus uncertainty (orientation, scale, and background pattern) and then compare human performance with that of the ideal observer, two biologically plausible model observers (simple MAX and normalized MAX). This experiment is complementary to an experiment we published earlier for a different subset of the five dimensions (Oluk & Geisler, 2023).

We find that our simulation method is quite efficient and can generate predictions for dense levels of variation along all five dimensions (and less dense variation along even more dimensions). We find that even after 5 million training images the CNN (ResNet18) approached but failed to reach ideal observer performance in our task, suggesting that our method can simulate ideal performance in naturalistic tasks that are complicated enough to be non-trivial for existing neural networks. We also find that normalizing the total energy of the templates of the MAX observer in addition to background normalization allows the MAX observer to approximate the optimal performance. Consistent with our earlier study, we find that the pattern of hits and correct rejections of the ideal, and normalized MAX observer (but not the simple MAX observer) are consistent with the human pattern. Comparison of human performance with and without the scale and orientation uncertainty (i.e., blocked scale and orientation conditions) showed a smaller difference than predicted by any of the models. However, we find that this smaller than expected difference can largely be explained by including biologically plausible levels of intrinsic position uncertainty in the ideal observer.

### Theory of Performance in High-Uncertainty Category Identification Tasks

Here, we define identification tasks in terms of a hierarchy of exhaustive and mutually exclusive events (Figure 2a). Events at the top layer of the hierarchy are object categories (*E*_*k*_), whose collection is the sample space (*Ω* = ⋃_∀*k*_ *E*_*k*_). For example, let event *E*_*k*_ represent the presence of the specific object k within the image displayed in a trial; *k* = 0 represents the event of no object. The task is to identify as accurately as possible which specific object k was presented in the trial (or if there was no object). In addition to trial-by-trial variability of object categories, most of the time in natural tasks, there are many other external sources of trial-by-trial variability, which we refer to as “extrinsic uncertainty.” In the category identification task, there are two kinds of extrinsic uncertainty: the variability in the properties of context (e.g., the contrast and spatial structure of the background) and the variability in the properties of object (e.g., 3D orientation and position of the object). In general, the properties of the context and of the object may be multidimensional. Here we will index the vector of properties of a specific context with the integer *l* and the vector of properties of a specific object with the integer *j*.

**Figure 2.**
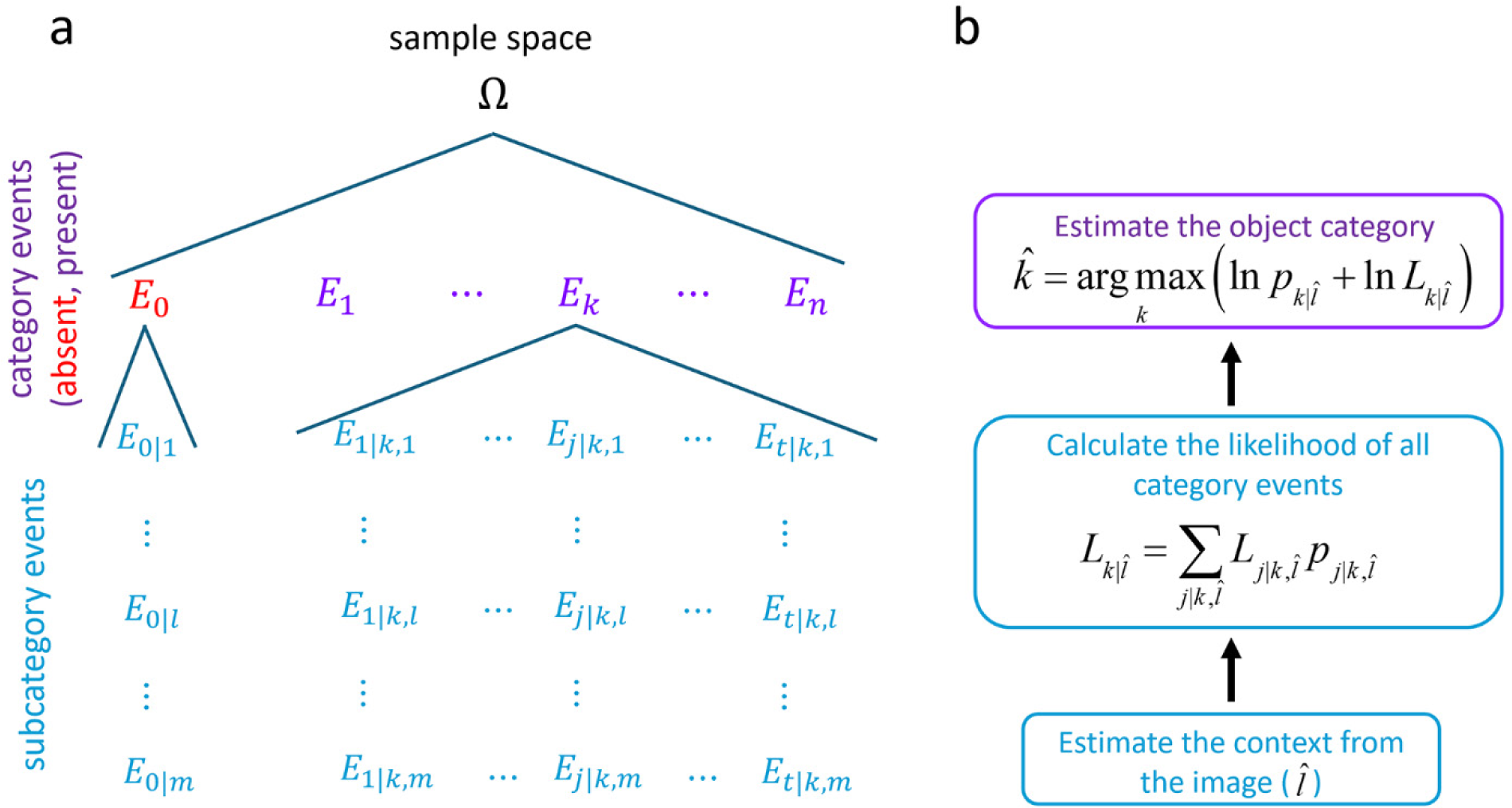
Target identification tasks under high levels of uncertainty represented as a hierarchy of exhaustive and mutually exclusive events. In each trial, the task is to identify the category event (k) based on the image presented. When there is an event of no object where the target is absent, it is possible to express it as a set of exhaustive and mutually exclusive subcategory events that cover the variations in the properties of background (extrinsic background uncertainty). Each target category event is also a set of exhaustive and mutually exclusive subcategory events that cover the variations in target properties in addition to background variations. We derived the optimal decision rule for any identification task expressed in this framework. First, we assumed that context variables can be estimated precisely from the image. The likelihood of each category is the sum of the subcategory event likelihoods multiplied by the prior probability of the corresponding subcategory event. Maximum a posterior probability estimate (MAP) is given by the maximum of the sum logarithm of category likelihoods with the logarithm prior probability of the corresponding category event. In sum, the optimal decision rule takes the maximum posterior probability of the category events, while the summation across posterior probabilities of subcategories gives each category event likelihood.

We incorporate the extrinsic uncertainties into the framework by describing each object category as a collection of multiple subcategory events constructing a bottom layer of the hierarchy. Specifically, the event of no object (*E*_0_) is composed of a set of subcategory events {*E*_0|*l*_} that are mutually exclusive and exhaustive of the event (*E*_0_ = ⋃_∀0|*l*_ *E*_0|*l*_). For example, when there is extrinsic uncertainty about background contrast, different levels of background contrast are indexed by *l*. Similarly, each object category (*E*_*k*_) is composed of a set of subcategory events {*E*_*j*|*k*,*l*_} that are mutually exclusive and exhaustive of the event (*E*_*k*_ = ⋃_∀*j*,*l*_ *E*_*j*|*k*,*l*_).

Here, we will focus on the identification of additive targets in white noise backgrounds. The only context variable associated with this environment is the standard deviation of the white noise background *σ*. Thus, the subcategory events of no target (*E*_0|*l*_) will reflect only variations in the standard deviation of the white noise background. On the other hand, subcategory events of target *k* (*E*_*j*|*k*,*l*_) will additionally reflect the set of targets associated with the subcategory (*a*_*j*|*k*_ **t**_*j*|*k*_). We define the target as the product of an amplitude term (*a*_*j*|*k*_) and a target profile whose Euclidean norm is one, ǁ**t**_*j*|*k*_ǁ = 1.

Note that in addition to target and background contrast variability, there is trial-by-trial variability of the exact white noise pattern. Thus, to be exact, any subcategory is composed of many subcategories corresponding to the variation in the exact white noise visual patterns. However, it is unnecessary to consider subcategories of white noise pattern because the relevant likelihoods can be computed directly from the image and target profile without having to separately consider each possible pattern. In other words, the primary reason for considering subcategories is to simplify to the point where it is possible to calculate the likelihoods from the image. One can stop at the point where the subcategories are simple enough to compute the likelihoods.

### Bayesian Ideal Observers

We have derived the optimal decision rule for the set of identification tasks, which can be characterized within the hierarchical structure described above (Figure 2a). The Bayesian ideal observer that maximizes overall accuracy chooses the object category *k* with the maximum posterior probability given the image presented on the trial *p*(*E*_*k*_|*I*). The probability of each category can also be written as a sum of probabilities of subcategory events *p*(*E*_*j*|*k*,*l*_|*I*) that are mutually exclusive and exhaustive sets of that object category. We define the likelihood ratio for each subcategory event (*L*_*j*|*k*,*l*_) to be the probability of the image given the corresponding subcategory event *p*(*I*|*E*_*j*|*k*,*l*_) divided by the probability of image given the subcategory of no object event with the same context index *p*(*I*|*E*_*j*|0,*l*_). In the Appendix (see the Derivation of the optimal decision rule), we derive the general version of the optimal decision rule:

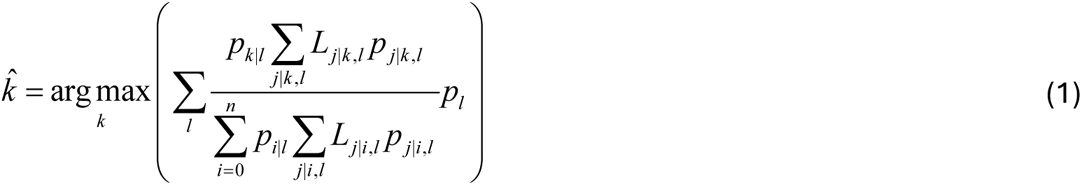

where *p*_*l*_ is the prior probability of the context given the image, *p*_*k*|*l*_ is the prior probability of category *k* given the context, and *p*_*j*|*k*,*l*_ is the prior probability of subcategory *j* given the category and context.

However, here, we only focus on cases where context variables can be precisely estimated, which makes it possible to simplify the optimal decision rule considerably (see Figure 2b). Given the estimated context variables *l̂*, the logarithm of the posterior probability of the object category can be expressed as a sum of the logarithm of the prior probability of the object category and the logarithm of the sum of the likelihood ratios of each subcategory scaled with prior probability of the subcategories:

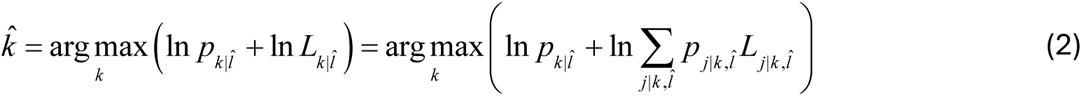

For the identification of additive targets in white noise backgrounds (as defined in the previous section), we can further simplify the general equation (see Appendix: Derivation of the optimal decision for additive targets in white noise). Specifically, we define the detectability (“d-prime”) for each sub-category event, 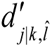, to be the target amplitude divided by the standard deviation of white noise *a*_*j*|*k*_⁄*σ̂*. We also defined the normalized template response as the dot product between image and the template divided by the standard deviation of the noise 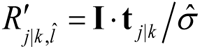. The following equation applies to any identification task involving additive targets in white noise:

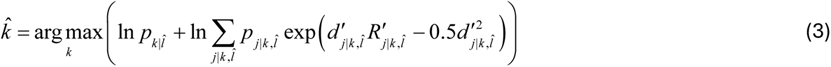

### Heuristic Observers

The most common heuristic for the identification tasks involving subcategories in any form (like position, time frames, etc.) is the substitution of the max operation for the summation over the likelihood ratios (Nolte & Jaarsma, 1967; Pelli, 1985; Graham et al., 1987). When the max operation operates over the ratio of likelihoods, its only advantage is avoiding the computationally intensive summation, thus enabling some further simplifications depending on the specifics of the task. In fact, by rearranging Equation 2 (see Appendix: Derivation of the heuristic max rule), it is possible to show that the max rule with likelihood ratios is the optimal decision rule if the task is to identify the subcategories:

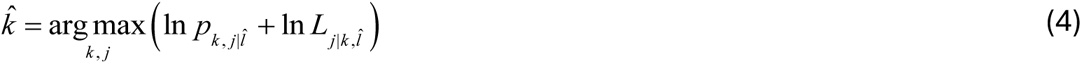

Here, we simplify the equation of the MAX heuristic observer for the identification of additive targets in white noise backgrounds (Equation 5). This parameterization of the max decision rule nicely separates the three potential sources of suboptimality: (i) the heuristic nature of the max operation, (ii) failures in the incorporation of the detectability map, and (iii) approximations to the normalized template response.

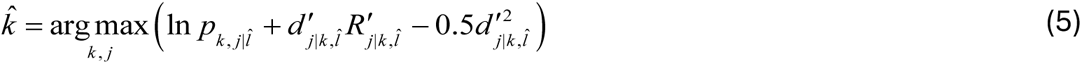

This MAX observer uses templates that are normalized to have unit energy and normalizes the template responses by the noise standard deviation. We will refer to this as the NN-MAX observer. In the current study with uniform prior probabilities, we use a simpler max rule over the normalized template responses together with arbitrary criteria *γγ* to approximate the ideal observer:

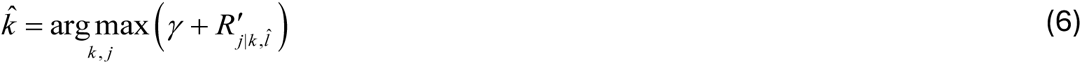

This observer also uses unit energy templates and normalized template responses but is simpler because it does not properly weight the normalized responses. We will refer to this observer as the SNN-MAX observer.

Finally, we consider an even simpler MAX observer with neither template nor template-response normalization. We will refer to this observer as the S-MAX observer.

### Convolutional Neural Network Observers

Recently, convolutional neural networks have reached human-level performance in many large identification tasks (He et al., 2015; Schmidhuber, 2015). CNNs are composed of many layers stacked on top of each other, making them deep neural networks. The central computations implemented in these layers are convolutions linked with the non-linear activation function for feature estimation and average/max pooling mechanisms, similar to computations utilized for optimal decision rules. However, it is often impossible to compare their performance to optimal performance because evaluating the ideal observer for large identification tasks is almost impossible due to the combinatorial explosion of the subcategories and non-well-behaved probability distributions. On the other hand, it is possible to compare these models for simple identification tasks (Reith & Wandell, 2020) to identify CNNs’ limits and weaknesses, which could help improve them. Here, we compare the performance of a CNN observer to that of the ideal observer for a more complex identification task. Moreover, since we trained the CNN from scratch, we could meaningfully examine the learning rate and how the relative performance across subcategories and contexts changes during the learning process. This information could potentially help to improve the learning process and network design.

For the current case of the identification of additive targets in white noise backgrounds, we decided to train the simple ResNet-18 architecture (He et al., 2016), which has been shown to approximate the ideal observer well under low levels of uncertainty (Reith & Wandell, 2020). ResNet-18 is composed of 18 layers of processing with skip connections and with the last stage a fully connected layer. Each layer involves convolution, batch normalization, and a non-linear operation (ReLu). Specifically, we used the implementation provided by Reith and Wandell (2020), in which the default Pytorch implementation (Paszke et al., 2017) was adjusted to perform a similar task. For example, the first layer was adjusted for gray-scale images, and the last layer was adjusted for the binary choices (for more details, see Reith & Wandell, 2020).

### Evaluating image-computable ideal and heuristic observers

Generating predictions of the ideal and heuristic observers becomes increasingly difficult as the number of conditions grows. The number of conditions (*d*) scales with the number of categories (*n*), subcategories (*t*), and context levels (*m*). As the *d* grows, the total number of trials that need to be simulated increases because ensuring there is at least a single trial for each condition is generally necessary for representative simulation. Moreover, in each of these trials, the subcategory likelihood ratios must be computed from the image. The number of subcategory likelihood ratios also scales with the total number of subcategories. Even though the subcategory likelihood ratio is a scalar number, its computation is repeated many times (it scales very rapidly with a number of conditions), making it computationally intensive because it involves images (large matrices). Repeating these image-related computations for each simulation experiment with different category-subcategory structures (uncertainty levels), prior probabilities, cost functions, etc., is incredibly expensive. Thus, we specifically aimed to develop a computational strategy that allows us to run many different simulation experiments while engaging the image-related computations only once.

Our strategy applies to any identification task with additive targets in white noise backgrounds and has the potential to generalize to different environments. We start by characterizing the set of target patterns the study concerns. In the present case, it is a set of raised-cosine-windowed sine waves with different orientations and scales. Then, we identify the computationally intense operations involving the images. Equation 3 shows that the subcategory likelihood ratios only depend on the dot products between templates and images, which can be decomposed into the dot product of two templates (one being the target presented in that particular trial) and the dot product of a background and template. We precompute the dot products of all possible pairings of template responses (all **t** *_j_*_|*k*_ · **t***_i_*_|*k*_) and the dot products of all possible templates with large set of randomly sampled backgrounds all having the same standard deviation, nominally 1.0 (all 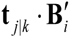). Once these template responses are stored, they can be used to evaluate the ideal and heuristic observers’ performance under various configurations of categories and subcategories as long as they only involve the pre-defined set of template patterns. We can evaluate the models under lower levels of extrinsic uncertainty and for various tasks (e.g., detection and discrimination). Moreover, we can change the prior probabilities, target amplitudes, and background contrasts associated with the simulation without computing any additional dot products. This greatly reduces the total computation time and allows running simulations with more stimulus dimensions and more levels per dimension. Demo code for MATLAB can be found on GitHub (github.com/CanOluk/Target_Identification).

### Simulations for High Levels of Target and Background Uncertainty

For the computational simulations, we implemented the equations and simulation strategy discussed in the previous section for the detection of additive targets in white noise backgrounds. As mentioned earlier, we simulated the effect on identification accuracy of target and background uncertainty for the five dimensions illustrated in Figure 1b. We chose these dimensions because they represent several of the important dimensions of variation that occur for target objects under natural conditions; specifically, random variations in 2D orientation, scale (distance from the observer), target amplitude/contrast, background contrast, and background spatial pattern. As mentioned earlier, likelihoods can be computed directly from the background spatial pattern and target template, so it was not necessary to consider subcategories of background spatial pattern. For the target, we picked a well-studied wavelet—a sinewave windowed with a raised cosine; its details are described in the experimental methods below. The target with a “scale-down factor” of 1.0 is defined as the largest target whose diameter is 3.35 visual degrees and spatial frequency is 1.0 cpd. (Here, a visual degree is 60 pixels.) Targets were scaled down by dividing the image coordinates by the desired scale-down factor. The scale-down factor increased linearly with the target’s spatial frequency, and consequently the target diameter decreases inversely with the spatial frequency. The scale-down factor was randomly sampled from eleven linearly spaced and equally likely values between 1 and 6. The orientation of the target was randomly sampled from 36 linearly spaced and equally likely target orientations between 0 and 360 degrees.

For the rest of the paper, we define the target amplitude as the single scaling factor, *A,* multiplying a target whose peak is normalized to 1.0. We decided on this definition of experimental amplitude because the changes in experimental amplitude correspond to the changes in the illumination of a fixed-reflectance pattern regardless of scale. However, with this definition of amplitude, the total energy of the target changes with the down-scaling factor. Nonetheless, for the simulations these experimental amplitudes can be easily transformed into the amplitude value, *a*, in the general equations by multiplying the experimental amplitudes with the total energy of the corresponding target. The experimental amplitude was sampled from sixteen linearly spaced levels, and the background contrast was sampled from eleven linearly spaced levels. We selected the range of background contrast and target amplitude so that the performance will rarely be perfect or at the chance level. For example, in the full uncertainty case, the proportion of hits for the smallest target (scale-down factor of 6) is 0.1 for the lowest amplitude and highest background contrast (hardest condition). In contrast, the proportion of hits in the easiest condition reached almost 1.0. In other words, we adjusted the computational resources to cover the dynamic range of performance. Moreover, these variations are still expected to have considerable effects in lower levels of uncertainty. For example, consider a low-uncertainty simulation (a blocked condition where none of these four dimensions vary) with the target scale-down factor of 4 (target 4 cpd, 0.85-degree diameter). For the ranges of variation of amplitude and background contrast in the high uncertainty simulation, the proportion correct in this low-uncertainty simulation ranges from 1.0 to 0.73.

We assumed that the prior probability distributions of target amplitude, scale, orientation, and background contrast are all uniform and that the probability of the target being present is 0.5. We evaluated the performance of model observers for various amplitude scalar factors, which is a single factor that multiplies the whole range of amplitude uncertainty. The decision criteria of the S-MAX and SNN-MAX observers were set to maximize the percentage correct.

Following our simulation strategy, we first precomputed the dot-products between the unit-energy templates and between the unit-energy templates and unit-variance white noise samples. A total of 396 templates were evaluated for 150,000 images, each of which was 201 x 201 pixels. The precomputation took less than two minutes with a reasonably powerful CPU (AMD Ryzen Threadripper 2950X 16-Core Processor 3.50 GHz). We determined psychometric function for the ideal observer by varying the amplitude scalar over ten levels, with almost 150,000 trials per level, which took around a minute. Thus, our strategy can handle large numbers of trials.

To compare various normalization computations that can be incorporated into the SNN-MAX observer, we also conducted another set of computational simulations, which revealed that our simulation strategy can handle many templates. A total of 18,360 templates are evaluated for sixty thousand 201×201 images (see Appendix: Figure A1), under the simultaneous target orientation (360 levels) and scale uncertainty (51 levels). Precomputation took around 10 minutes. Evaluating a psychometric function for the ideal observer for thirty levels of amplitude scalar, each, including sixty thousand trials, only took about 2 minutes. Note that this is equivalent to simulating 4 dimensions of uncertainty with 10 levels along each dimension.

### Simulation results: Bayes ideal and max observers

First, we simulated optimal performance for lower levels of uncertainty to compare with the high uncertainty case. For the lowest level of uncertainty, performance is the average across blocked conditions that provide no extra uncertainty (black diamonds in Figure 3). As extrinsic uncertainty increases, the task becomes more difficult, as indicated by the lower optimal performance. The light-blue diamonds and light-green circles show the performance of the ideal and SNN-MAX observers with target amplitude and background contrast uncertainty. The mid-blue diamonds and mid-green circles show the performance of the ideal and SNN-MAX observers with target orientation and scale uncertainty. The dark-blue diamonds and dark-green circles show the performance of the ideal and SNN-MAX observers for all dimensions of uncertainty. The relatively big drop in performance when orientation and scale uncertainty are included is due to the fact that the background contrast can be precisely estimated from the image presented in each trial. Figure 3 also shows that the SNN-MAX observer performs almost as well as the ideal observer (Equation 3).

**Figure 3.**
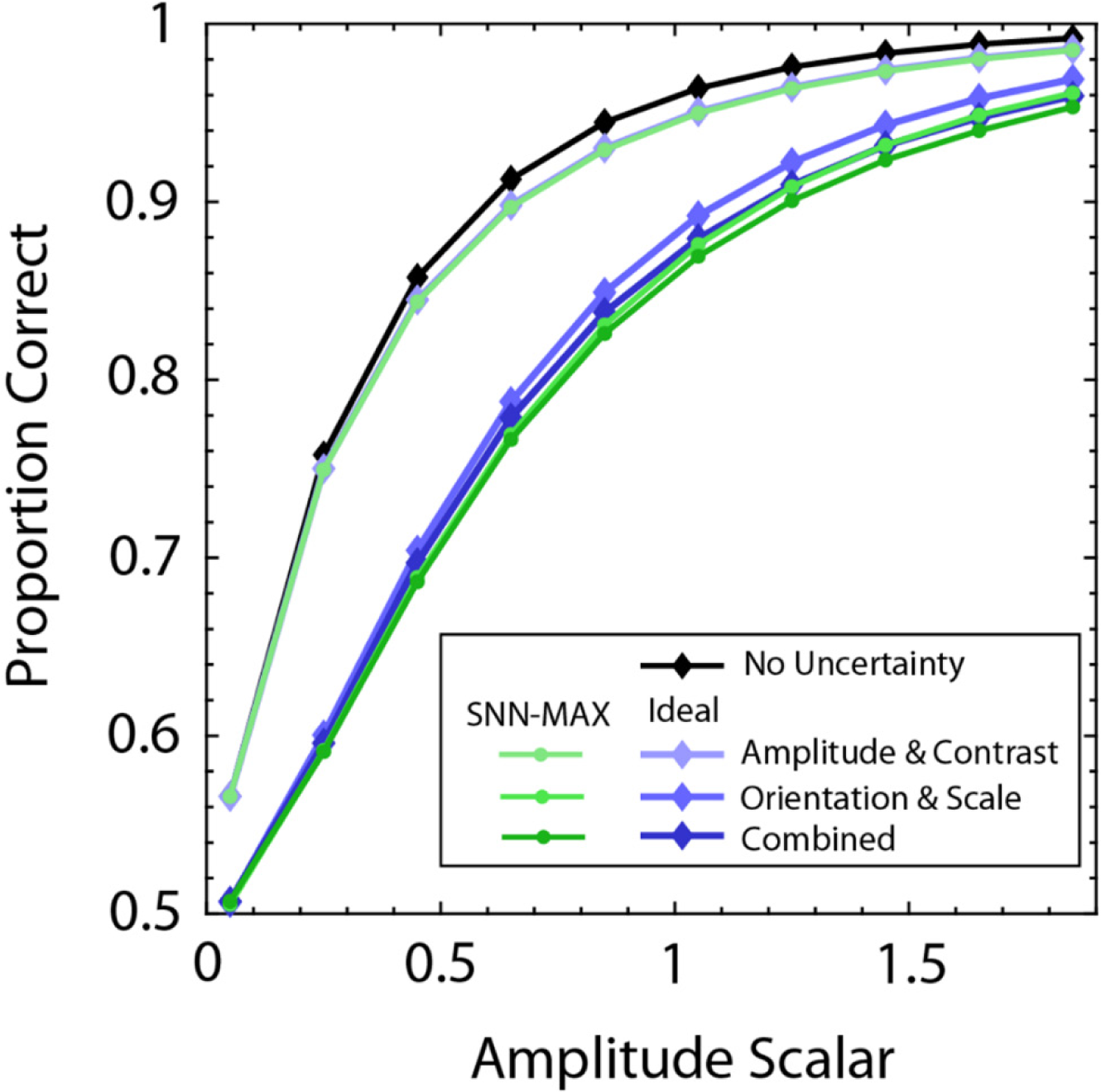
Computational simulations for the ideal and normalized MAX observers. The ideal and normalized MAX observers’ performances are shown in blue and green, respectively. In the x-axis, we plotted an amplitude scalar, a multiplicative scalar factor on the range of amplitudes used to generate the target amplitude uncertainty. In the combined condition, target scale, orientation, amplitude, and background contrast are all randomly sampled. There are 36 orientations, 11 scales, 16 amplitudes, and 11 background contrast levels, adding to a total of 69.696 different target-present conditions. We simulated a psychometric function containing ten amplitude scalar levels (for each level, there were 139.392 trials) with a novel method we developed based on pre-computations. In this case, the pre-computation took about two minutes. The psychometric function for the ideal observer under combined uncertainty is generated in a minute. (Processor: AMD Ryzen Threadripper 2950X 16-Core Processor 3.50 GHz). The performance is shown with lighter colors for lower uncertainty levels, illustrating the effect of the prior information. We have specifically simulated the performance under simultaneous amplitude and contrast uncertainty (similar to Oluk & Geisler, 2023); and simultaneous scale and orientation uncertainty (will be studied below). When a level of a certain dimension is fixed in a block of experiments, it provides prior information about the dimension, so models benefit from it. The performance shown here is averaged across dimensions that do not vary trial by trial. Also note that when there is no uncertainty about these four dimensions, both models make the same predictions.

We also simulated other heuristic model observers to compare with the ideal and SNN-MAX observers. For the SNN-MAX observer, in separate simulations (see Appendix: Figure A1), we focused on orientation and scale uncertainty with a much denser sampling of each dimension to test various normalization methods for templates. We found that using non-normalized templates with equal peaks of 1.0 performed the worst (S-MAX observer), while using templates with equal energy (the SNN-MAX observer) performed the best.

Previously, we showed that the S-MAX observer performs much worse than the SNN-MAX observer under the simultaneous target amplitude and background contrast uncertainty (Oluk C Geisler, 2023). On the other hand, we find here that the NN-MAX observer (Equation 5) performs only very slightly better than as the SNN-MAX observer (Equation 6), even when there is simultaneous uncertainty about the target amplitude, orientation, scale, and background contrast (the combined conditions in Figure 3; see Appendix: Figure A2). This result is consistent with our recent work showing that incorporating the exact detectability map or prior probability map is not necessary to approximate the optimal decision rule in visual search tasks, where the task is to identify target absence or the target location if present (Zhang C Geisler, 2024). Together, all these results suggest that both normalizations (background contrast and template energy) are necessary for the MAX observer to approximate the optimal performance, whereas incorporating the exact detectability map is not necessary.

### Simulation Results: Resnet18

We also evaluated the performance of a convolutional neural network for the full uncertainty case. Specifically, we picked the amplitude scalar level of 1.05, where the optimal performance reaches almost 88 percent correct. We trained from scratch the ResNet-18 architecture that was previously used to approximate optimal performance with low levels of extrinsic uncertainty (Reith & Wandell, 2020). We trained the neural network with 5 million unique images while regularly testing the performance over a separate set of seventy-five thousand test trials. The performance in the training and test data steadily improved without any sign of overfitting, likely due to the training images being new each time (Figure 4A, black and red curves). The results show that the convolutional neural network achieves excellent performance (0.8) very quickly. However, it reached the general normalized SNN-MAX observer’s performance after 2.5 million images and failed to reach the optimal performance with 5 million images. Thus, we can effectively simulate the ideal observer and SNN-MAX observer for tasks where it is not trivial for a convolutional neural network to achieve the same level of performance.

**Figure 4.**
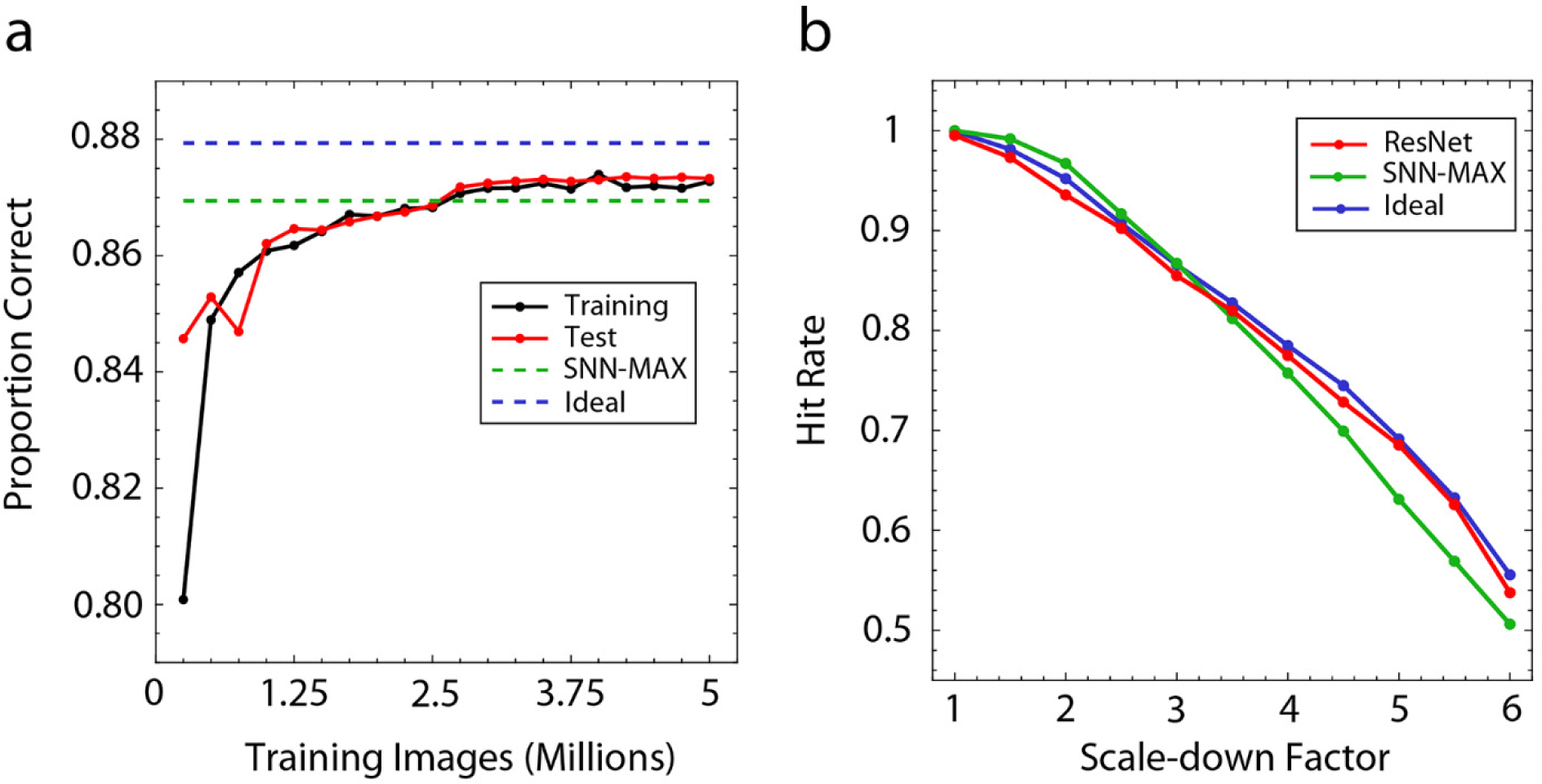
For the amplitude scalar of 1.05, we have trained a ResNet-18 with 5 million training images. In the left panel, ResNet-18 performance is shown as a function of the number of training images. The black line corresponds to the performance evaluated in the training dataset so far, and the red line indicates its performance in the test dataset of seventy-five thousand images. Approximately after 2.5 million images, ResNet’s performance reaches the normalized MAX observer’s performance (0.869). The performance of ResNet kept increasing shallowly but failed to reach the ideal observer performance (0.879). Finally, we evaluated the trained ResNet with the same dataset used for other models, and its performance was found to be 0.873. In the right panel, the hit rate as a function of the target scale is shown for all models evaluated with the same dataset. Specifically, the normalized MAX observer’s hit rate drops sharper than the ideal observer’s. However, ResNet approximates the ideal observer’s hit rates with a similar slope.

There are a couple of advantages to studying relatively difficult tasks where it is possible to compare machine learning and ideal observer methods in a non-trivial manner. First, it can facilitate our understanding of the computational mechanism of these visual processes because it is possible that they may make different predictions that can be tested even though here, we have not further tested ResNet-18 against the data due to its similarity to the ideal observer. Secondly, benchmarking the neural network performance against the optimal performance and simple heuristics can help us better understand the task’s computational requirements. For example, to understand whether the trained network is systematically different in certain conditions, we plot the hit rate of the model observers as a function of the scale-down factor (Figure 4B). We see that Resnet-18 seems slightly and almost equally worse in all conditions, so the error is not systematic, which we find also holds for different dimensions of uncertainty.

### Human Performance Under High Levels of Orientation and Scale Uncertainty

Under natural conditions, the visual detection task involves a vast amount of extrinsic target and background uncertainty along multiple stimulus dimensions. Unlike natural conditions, laboratory experiments are often conducted under strictly controlled conditions with relatively little extrinsic uncertainty. However, conducting experiments under more naturalistic conditions with higher levels of uncertainty is important for getting a comprehensive understanding of the visual mechanisms underlying target identification. For example, a visual target’s 3D orientation and position relative to an observer vary enormously. Here, we measured human ability to detect additive targets in white noise when 2D target orientation and target scale randomly vary over a wide range. Computer simulations (see the previous section) allowed us to identify three candidate models: the ideal observer, SNN-MAX observer, and S-MAX observer. In our previous study (Oluk & Geisler, 2023), we showed that the hit and correct rejection rates can be used to distinguish between model observers under high levels of uncertainty. We found that these models (see Appendix: Figure A3) make different predictions for the pattern of hit and correct rejection rates. Thus, we compared these three models against the human data.

We also measured human detection performance under low uncertainty when target orientation and scale were fixed in a single block. The orientation and scale levels were the same as those used in the high-uncertainty experiment. Based on the performance under low uncertainty, the decrease in the overall performance due to the introduction of high levels of uncertainty can be predicted. We aimed to test these theoretical predictions by comparing human efficiency across the high and low uncertainty conditions (Cohn & Lasley, 1974; Swensson & Judy, 1981; Davis et al., 1983; Cohn & Wardlaw, 1985; Burgess,1985). Previous studies focused on the effect of uncertainty of each of these dimensions in isolation. Orientation uncertainty in isolation has been found to have either no or minimal effect on human performance (Doehrman, 1974; Ukkonen et al., 1995). However, these measurements were not compared to theoretical predictions. There are no published studies on scale uncertainty because when the target’s scale changes (for example, when the target’s position in-depth changes), both the target’s spatial frequency and size change in relation to each other. Human performance has been measured for target spatial frequency uncertainty and target size uncertainty while keeping one of the two components fixed. Moreover, in these studies, the discriminability of different target levels (sizes or spatial frequencies) was equated by adjusting amplitudes for different targets. They found that the effect of size uncertainty in isolation is smaller than theoretically expected based on the data measured under low extrinsic uncertainty (Judy et al., 1995; Judy et al., 1997). In contrast, the effect of spatial frequency uncertainty in isolation is roughly consistent with the expected effect, even though results are somewhat inconsistent across studies (Davis & Graham, 1981; Davis et al., 1983; Yager et al., 1984; Hübner, 1996a; 1996b; Ohtani et al.,2002). Thus, studying the effect of simultaneous target orientation and scale uncertainty on human performance should provide new insights into how the brain utilizes the prior information. Moreover, these results can help us to differentiate the effect of multiple computational mechanisms such as attentional bottlenecks (Lu C Dosher, 1998; Lee et al., 1999), intrinsic uncertainties, and reductions of uncertainty (Tanner, 1961; Pelli, 1985).

### Experimental Methods

The target was a sine wave windowed with a radial raised-cosine function. The background was a square patch of white noise with a width of 12 visual degrees (721 pixels). The target was presented at the center of the display, together with the white noise background. The range of target orientations was the same as in the simulations (0 to 359 degrees) but was sampled with eight levels. The range of scale-down factors was smaller than in the simulations (1 to 4.5 instead of 6) and was sampled with eight levels. The eight linearly spaced levels of target orientations were 0,45,90,135,180,225, 270, and 315 degrees. The eight linearly spaced levels of scale-down factor were 1,1.5,2,2.5,3,3.5, 4, and 4.5 (Figure 5). The target amplitude and background contrast were selected to give an accuracy of around 75 percent correct. The target amplitude (*A*) was 3.5 gray levels, and the standard deviation of white noise was 15.46 gray levels (12 percent RMS contrast). The background gray level was 128 (47.18 cd/m2)

**Figure 5.**
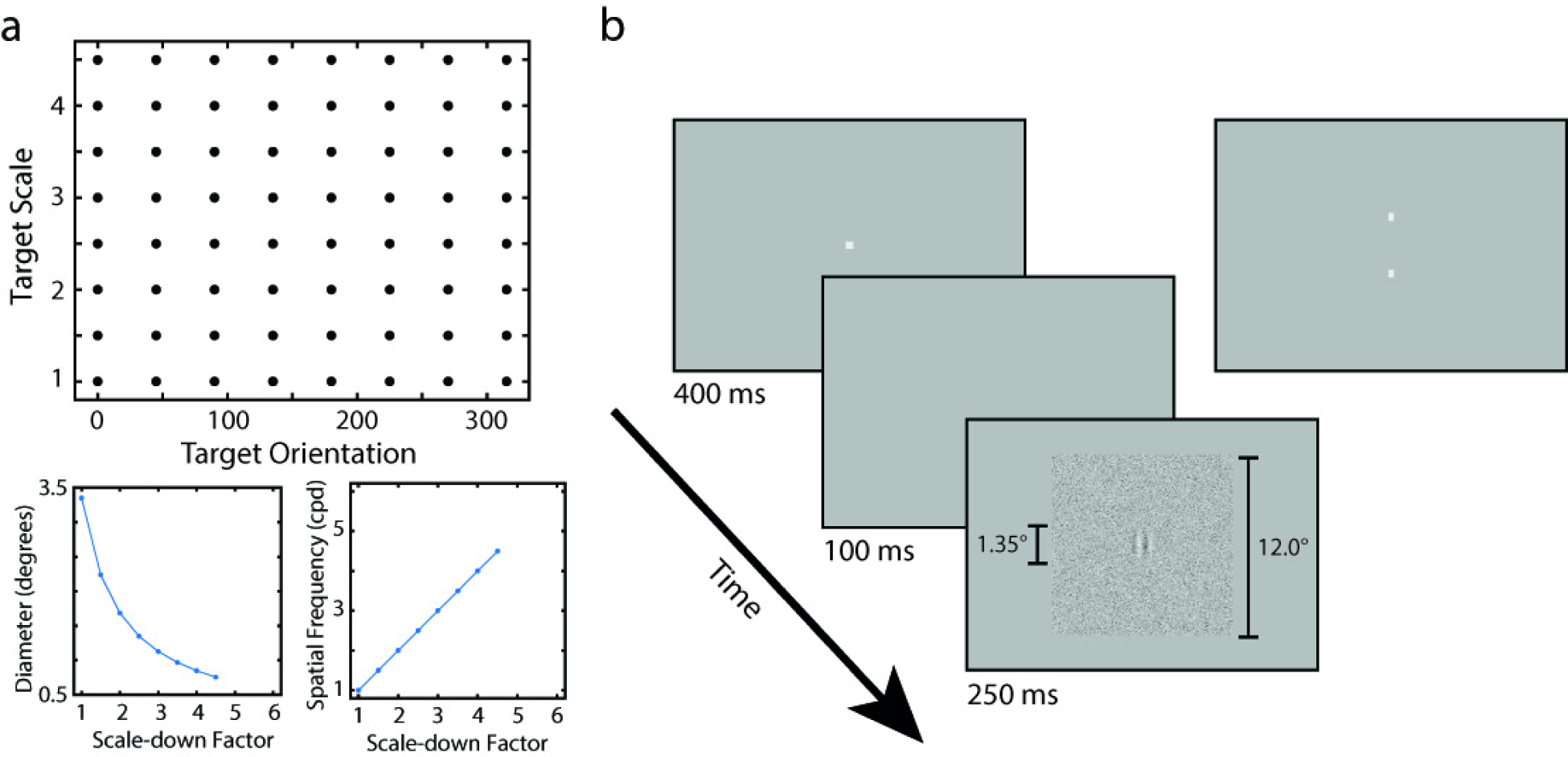
**a** Target scale and orientation levels in the experiment. For any target present trial, the stimulus’s parameters are uniformly sampled from the grid of points. The two small figures show the relationship between the target scale and the target’s diameter and spatial frequency. **b** The procedure of the experiment. 400 ms fixation screen is followed by a 100 ms black screen. The stimulus is shown for 250 ms with twelve visual degrees square background. In the low-uncertainty condition, the procedure was the same except for the fixation shown on the side. The fixation consists of two rectangles similarly oriented to the target and always 20 percent of the target diameter away from the target location to remind participants of the scale and orientation of the target presented in that block. The example shown here has an orientation of 90 degrees and a scale of 2.5.

Participants were asked to press a key in each trial to indicate whether the target was present or absent. Each trial started with a dim fixation point at the center, presented for 400 ms, followed by a 100 ms blank background. The stimulus was presented for 250 ms (Figure 5). Participants had 750 ms to indicate their decision, and the subsequent trial started after an additional 700 ms. The stimulus presentation time was selected to match the average fixation duration of the human eye during covert visual search. Audio feedback was provided to participants in each trial.

On each trial, the target scale and the orientation were randomly and independently sampled from uniform distributions over eight discrete levels of scale and orientation; however, the sampling was counterbalanced, so there were the same number of trials for each condition. The biggest and smallest target (at high amplitude) was shown on screen in the breaks to aid participants in remembering the range of scales that could be presented in the experiment.

For the low extrinsic uncertainty experiment, there were 64 different blocks of the experiment; at the start of each block, the target for that block (at high amplitude) was shown to make sure that participants knew which target was to be presented in that block. The fixation target also depended on the target (Figure 5). The fixation target was composed of two small rectangles that were separated from the target location by 20 percent of the target’s diameter and were also oriented parallel to the orientation of the bars of the sinewave.

The experiment consisted of two sessions. Each session was a complete experiment with only half of the trials. In each session, participants first completed the low uncertainty condition that contains 64 blocks (different target conditions), each one with 26 trials (13 target present, a total of 52 trials with two sessions). The order of blocks was randomized. The high uncertainty condition is split into five blocks, the first three completed in the first session and the last two completed in the second session. Each high uncertainty block consists of 640 trials for a total of 3200 trials.

Three experienced and one naive observer completed the experiment. They had normal (corrected) spatial acuity. Written, informed consent was obtained for all observers in accordance with The University of Texas at Austin Institutional Review Board. The stimuli were presented at a distance of 90 cm with 60 pixels per degree resolution. The luminance of the background was the mean of the display range (47.18 cd m2). Images were gamma-corrected based on the calibration of the display device (Sony GDM-FW900), quantized to 256 gray levels. The screen refresh rate was 85 hertz. All experiments and analyses were done using custom codes written in MATLAB with the Psychophysics Toolbox (Brainard, 1997; Pelli, 1997).

### Analysis Methods

The mean gray level was subtracted from images for computational simulations. Simulations were made with the target scale and orientation levels the same as those used in the psychophysical experiment. The three model observers were fitted to the data with the same likelihood method introduced in the previous study (Oluk & Geisler, 2023). The method implicitly fits the hits and correct rejection rate of the subjects to maximize the likelihood of the data by varying two parameters. The first parameter is the scale factor on the target amplitude. The second parameter is a criterion. Here, we made a slight adjustment to the likelihood method. Sometimes, due to a finite number of simulated trials, the model achieves a hundred or zero percent correct, making the likelihood of the data zero. To avoid the issue, we assumed there would be a single error if we doubled the number of simulated trials. We calculated rates based on this assumption. If target orientation and scale are known, all three model observers perform identically and are optimal. For 64 blocked conditions in the low uncertainty condition, only a single overall scalar on amplitude was varied in the fitting procedure. Criteria were calculated from the data and fixed. The rest of the fitting procedure was the same as for the high uncertainty condition. The data from both conditions were fitted simultaneously to test principles about how the low-uncertainty condition is related to the high-uncertainty condition.

Intrinsic position uncertainties were simulated by including additional potential visual targets whose centers vary around the actual center of the presented target. The maximum position uncertainty is 12 pixels in radius, resulting in a 25 x 25 pixel square that involves all possible center locations centered around the true center. However, introducing 624 new locations is computationally expensive (for each new template location, there are 64 templates due to orientation and scale uncertainty, so the total number of templates would be 39,936) and unnecessary because these templates are highly correlated. Thus, new templates were generated for two-pixel steps, effectively creating 13 x 13 possible center locations. Because of that, the amount of position uncertainty is starting from 2 pixels to 12 pixels with steps of 2. However, we found that this type of position uncertainty has little effect on large templates because only a tiny percentage of templates are sampling new spatial locations. Thus, we simulated another hypothesis, which is that intrinsic position uncertainty might be fixed in terms of the number of neurons. Neurons with larger receptive fields also tile the visual space, but their centers are more separated. Thus, for neurons with larger receptive fields, the same amount of position uncertainty would correspond to larger shifts in the center of the target. In this case, the implementation of position uncertainty stays the same as for the smallest target. However, for any other targets, steps (and radius) are scaled up by the smallest scale divided by its scale (for example, the largest target has a scale of 1, so a 12-pixel radius multiplied by 4.5 and rounded to the 54-pixel radius, and steps are 9 pixels steps).

## Results

The average observer’s data was obtained by aggregating the data over participants. Confidence intervals were generated by resampling the data, assuming the response in a trial follows a binomial distribution with a success rate of corresponding hit (miss) and correct rejection (false alarm) rates. Confidence intervals for fitted parameters were computed by fitting the bootstrapped data.

The hit rates for the 64 conditions and the correct rejection rate of the average participant in the uncertainty experiment are shown in the left panel of Figure 6. We found that the ideal observer (negative log likelihood with 95 percent confidence intervals: 14377 [14178 −14561]) and SNN-MAX (14602 [14405 −14787]) better explain the data than S-MAX (15461 [15162 −15748]). Both the ideal and SNN-MAX observers explain most of the variation in the hit (RMSE: 0.07 and 0.09) and correct rejection rate (RMSE: 0.01and 0.03) whereas S-MAX failed to do the same for hit (RMSE:0.13) and correct rejection rate (RMSE:0.08). (See Appendix: Figure A4 for best-fitted predictions of the models.) The estimated scalar factors on amplitude are similar across all observer models (Ideal: 0.41, SNN-MAX: 0.38, S-MAX: 0.56). The similarity of the estimated scalar factors across models means that the overall performance of model observers is similar (with S-MAX being slightly worse). The estimated criterion, when converted to the prior probability of the target-absent that it would be optimal for, is also similar for all observer models (Ideal: 0.45, SNN-MAX: 0.46, S-MAX: 0.47). The similarity of the estimated criterion across models means all three models agree that the human observers are biased slightly toward “target present.” Both parameters account for suboptimalities of human participants regardless of the model. Thus, differences in the quality of fit are more about the pattern of errors across the stimulus dimensions than overall performance or bias.

**Figure 6.**
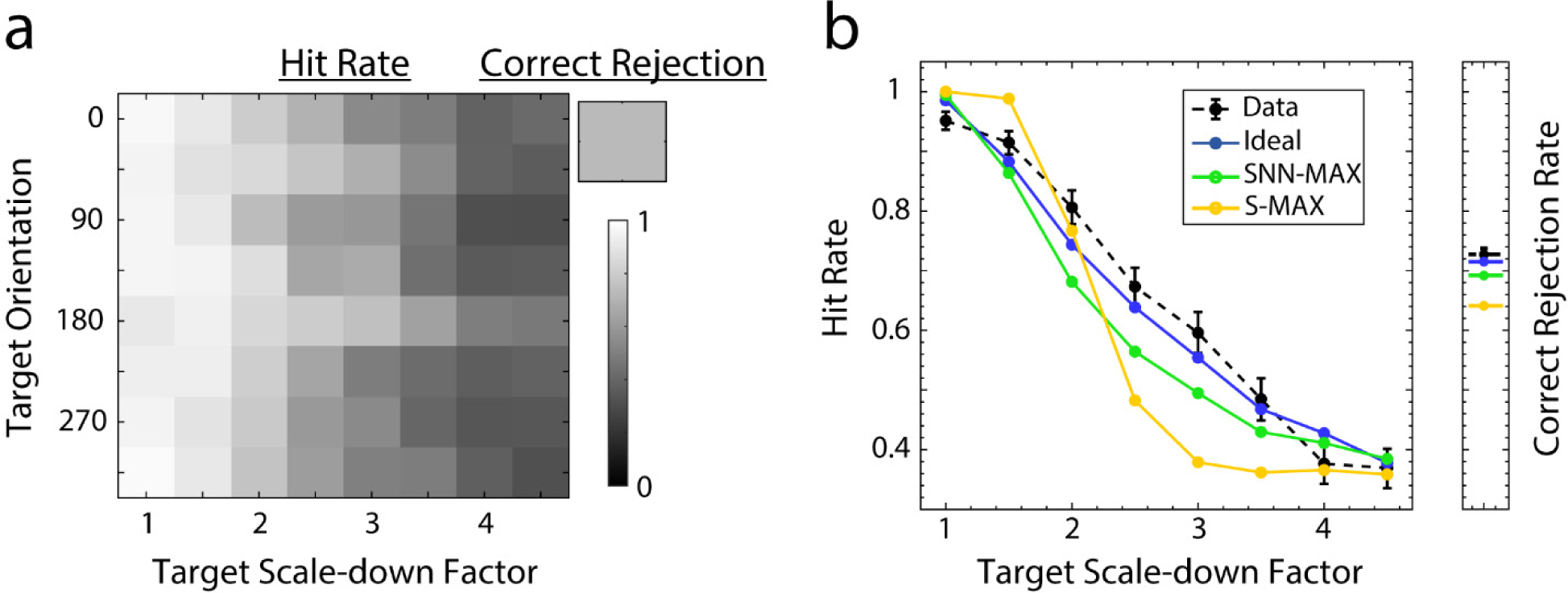
**a** We measured the human detection performance under a simultaneous target scale and orientation uncertainty. The overall percentage correct of the average participant is 0.69. Hit and correct rejection rates of the average observer are shown for each condition (sixty-four different target-present, one target-absent condition). We found that there are no large consistent performance differences between different orientations. **b** We have fitted the ideal, normalized MAX, and simple MAX observer whose templates are not normalized (thus have different total energies) to the average participant’s hit and correct rejection rates. Specifically, we varied a criterion parameter and overall scale factor on amplitude to maximize the likelihood of the data. The overall percentage correct of fitted ideal, normalized MAX, and simple MAX are 0.70, 0.65, and 0.61, respectively. The average participant’s hit rate as a function of scale (averaged across orientations) is shown together with model observers’ best-fitted predictions. Error bars around the data show 95 percent confidence intervals.

The pattern of results is robust across participants (see Appendix: Figure A5). For two participants (P2 and P3), the data is better explained by the ideal and SNN-MAX than by the S-MAX model. For the other two participants (P1 and P4), the 95 percent confidence intervals around the negative log-likelihoods overlap, but there is still a clear trend in the same direction. The quality of the fits is best illustrated with the correct rejections and hit rates averaged across the orientation levels because there is little variation with the orientation. For the average observer, even the best-fitted S-MAX observer predicts a much lower correct rejection rate and a sharper decline in hit rates as a function of the target scale (Figure 6B). Even though the ideal and SNN-MAX are better at capturing the pattern, they also fail systematically by predicting a slightly sharper decrease as a function of scale (compared to the approximately linear decrease of data) and a slightly lower correct rejection rate.

We have fitted the ideal observer to the average data collected in the low uncertainty condition. It explained the most variation in the data (R^2^ for hit rate matrix: 0.86, R^2^ for correct rejection matrix: 0.36) with the estimated amplitude scale factor of 0.3 (see Appendix: Figure A6). If the inefficiency of participants is due to simple internal noise, one would expect the same scale factor to translate to the high uncertainty condition. We generated predictions of the ideal observer for the high uncertainty condition based on the scale factor estimated from the low uncertainty condition and the optimal criteria (Figure 7A, blue points). The prediction is quite poor, and the human performance is much better in the high uncertainty condition than the predicted performance based on the low uncertainty condition.

**Figure 7.**
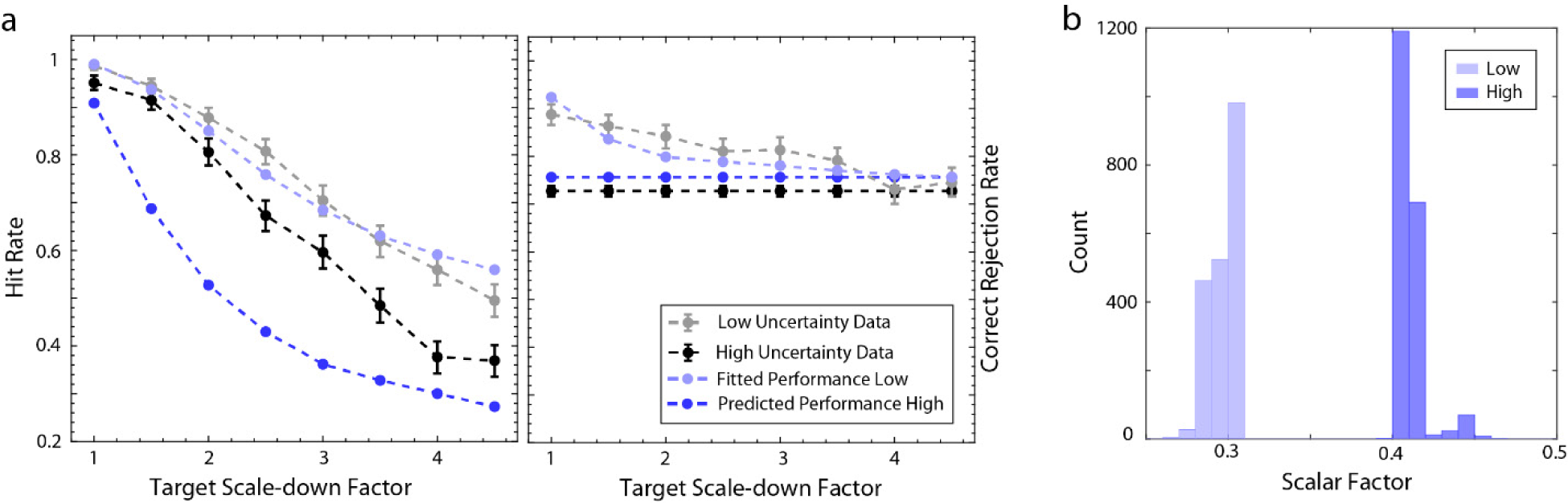
**a** We also measured the human performance in the low uncertainty condition where both scale and orientation are fixed in a block of the experiment. The percentage correct of the average participant is 0.78. Average participant’s hit rates as a function of scale are shown with a dashed gray line under low uncertainty and with a dashed black line under high uncertainty. The ideal observer for the low uncertainty is fitted to the average observer’s hit and correct rejection rates by only varying a single scalar factor on the amplitude. The best-fitted model’s overall percentage correct is 0.78, and its predictions as a function of scale are shown with a solid light blue line. The estimated scale factor under low uncertainty is used to generate predictions for the high-uncertainty condition with optimal criteria shown with a solid dark blue line. The predicted overall percentage correct is 0.62, whereas the average participant’s percentage correct is 0.69. Error bars around the data show 95 percent confidence intervals. **b** The distribution of scale factors is estimated by fitting the ideal observer to the low and high uncertainty conditions separately for two thousand bootstrapped versions of the average participant’s data.

Next, we directly compared the estimated scale factors by fitting the ideal observer to either the high uncertainty condition or the low uncertainty condition in isolation. If the efficiency of human observers was the same for both conditions, we expect the estimated scale factors to be the same, consistent with the parallel processing of the information (no further limitation for higher uncertainty conditions). If there were limited attentional capacity, the performance would decrease more than predicted because many more patterns would need to be attended to/tracked in the high uncertainty condition. Thus, limited attentional capacity (and various other computational limitations that are only effective for high uncertainty conditions) predict a lower scalar factor in the high uncertainty condition because they predict lower efficiency under high uncertainty. The distribution of estimated scale factors was computed by fitting the bootstrapped average participant data. The estimated scale factors for the high uncertainty experiment are higher than for the low-uncertainty experiment (Figure 7b). Therefore, the visual system is more efficient in high uncertainty conditions, suggesting that neither the uncertainty reduction hypothesis nor attention bottleneck is sufficient to explain the data alone. Some of the inefficiency in the low uncertainty condition must not translate into the high uncertainty condition.

An important factor that could explain the relationship between low and high uncertainty conditions is the existence of intrinsic uncertainties (Tanner, 1961; Pelli, 1985). Even when there is absolutely no extrinsic uncertainty about the target (for example, if target orientation, scale, and position are fixed and known), the visual system cannot exploit this information perfectly and must consider multiple alternatives about the properties of the target and background while making an identification decision. Such intrinsic uncertainty would have a considerable effect when there is low extrinsic uncertainty but almost no effect when there are high levels of extrinsic uncertainty. To test whether the combined effect of orientation and scale intrinsic uncertainty is a plausible explanation for our results, we simulated model observers with extremely high levels of intrinsic uncertainty (+− 10 degrees of orientation and +− 1 level of the scale-down factor). When the ideal observer’s efficiency is reduced to match human performance in the low uncertainty condition, the predictions of the ideal observer with intrinsic uncertainty still falls short of human performance in the high uncertainty condition. Therefore, these two dimensions of intrinsic uncertainty cannot solely explain the observed difference between the low and high extrinsic uncertainty conditions.

Intrinsic position uncertainty has a bigger effect than orientation and scale uncertainty on the overall percentage correct. For the scaled version of position uncertainty (uncertainty is proportional to target size; see Methods for details), with a plausible level of intrinsic position uncertainty (0.1 visual degrees radius for smallest target), the predictions are consistent with the overall difference in human accuracy between the low and high uncertainty condition.

Next, we tested whether the model observers can account for the pattern of hits and correct rejections. The model observer with intrinsic position uncertainty was fitted to data of both conditions simultaneously while varying three parameters: (i) the decision criterion for the high uncertainty condition, (ii) a single amplitude scale factor for both conditions, and (iii) the level of intrinsic uncertainty. The model without intrinsic uncertainty was fitted simultaneously to both conditions, providing a baseline for comparisons (dashed blue curves in Figure 8). Our analysis revealed that the model with intrinsic position uncertainty (solid blue curves) did not better explain the data in terms of negative log-likelihood compared to the baseline. However, when the difference between the average participant’s and the fitted model’s hit and correct rejection rates was quantified with root-mean-squared error (RMSE), the model observers with intrinsic uncertainty did better explain the data. The reason behind the difference between RMSE and negative log-likelihood measures can be appreciated by examining Figure 8. The model with intrinsic position uncertainty better explains the overall performance difference between the high-uncertainty and low-uncertainty conditions, which is much smaller than the baseline model predicts. However, the model with intrinsic uncertainty predicts a sharper fall with the target scale than the human data, which is better captured by the baseline model (especially in low-uncertainty conditions). Because of the sharp decrease, the model with intrinsic position uncertainty produces some hit and correct rejection rates close to a hundred percent, producing very large negative log-likelihood errors.

**Figure 8.**
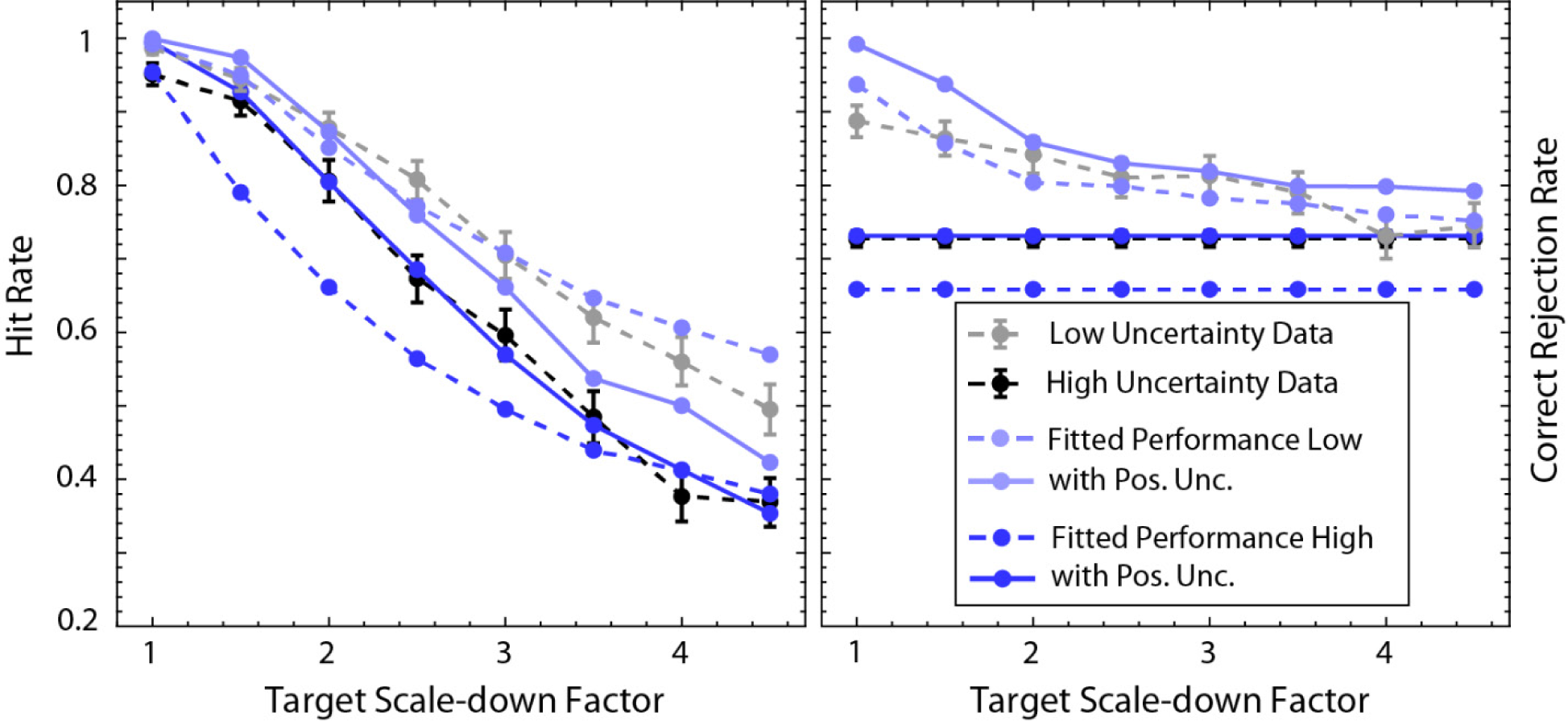
The ideal observer with and without intrinsic position uncertainty is fitted to both low and high uncertainty conditions simultaneously by varying a single scalar factor on amplitude and a criterion for high uncertainty condition. The overall percentage corrects of the average participant, and best-fitted ideal observer without intrinsic uncertainty are 0.78, 0.81 (low uncertainty), and 0.69, 0.63 (high uncertainty), respectively. On the other hand, the ideal observer with a scaled intrinsic position uncertainty of 0.13 visual degrees radius (for the smallest target) predicted 0.78 and 0.69 overall percent correct for low and high uncertainty conditions, respectively. The predicted hit rates (averaged across orientation) and correct rejection rate as a function of scale are shown as solid blue lines for the ideal observer with intrinsic position uncertainty.

## Discussion

We measured the effect of simultaneous extrinsic target scale and orientation uncertainty on human ability to detect wavelet targets in white noise backgrounds. As expected, the uncertainty had a substantial effect on overall accuracy. The hit rate declined substantially as a function of the scale-down factor for both the high and low extrinsic uncertainty conditions but was relatively constant as a function of orientation (black and gray symbols in Figures 6, 7 and 8). The pattern of results is not consistent with an ideal observer limited by only the extrinsic uncertainty and background noise and is more consistent with an ideal observer (or SNN-MAX observer) that is also limited by plausible levels intrinsic position uncertainty.

As can be seen in Figures 6-8, performance declined monotonically with the scale-down factor of the target. This is consistent with previous measurements with fixed bandwidth wavelets on uniform backgrounds (Watson, 1987; Watson & Ahumada, 2005). Also, we found that performance was relatively flat as a function of target orientation. In other words, we did not observe a substantial oblique effect. This is also not surprising. When targets appear on uniform backgrounds, detection and discrimination thresholds tend to be maximum when the target orientation is oblique (Appelle, 1972), consistent with the high density of cortical neurons tuned to the cardinal directions (Mansfield, 1974; Mansfield & Ronner, 1978; De Valois et al., 1982; Coppola, White, Fitzpatrick, & Purves, 1998; Furmanski & Engel, 2000). On the other hand, in high contrast naturalistic noise, detection of oriented structure is best for the oblique directions (Essock et al. 2003; Hansen & Essock 2004). As these authors suggest, this could be due orientation-tuned gain control (normalization), which is stronger in the cardinal directions because of the greater numbers of neurons contributing to the gain-control signal. With white noise backgrounds, and depending on the noise contrast, it is plausible to get anything in between, including little or no oblique effect, which we observed.

We also found that normalizing the energy of templates is necessary for the max observer to account for the human data and for the max observer to approximate the ideal observer. Energy normalization ensures that the inputs provided to the max operation have equal variance. Thus, no subset of input channels with produce larger random responses to the noise that will tend to dominate the max operation. We consider two plausible neural implementations of energy normalization that make testable predictions. First, it is possible that neurons with different sizes of receptive fields are wired to receive a similar amount of excitatory and inhibitory inputs in total energy. These wiring patterns likely result in similar levels of noise variance, which should be testable either without any visual input or with white noise input. Secondly, it is also possible to implement normalization in the upstream neurons responsible for implementing the max operation. These neurons might weight each input by its baseline variance, which might also be testable.

We found that human efficiency relative the ideal observer is higher under high orientation and scale uncertainty than under low uncertainty, and that this could be explained by plausible levels intrinsic position uncertainty (see Figure 8). Plausible levels of intrinsic scale and orientation uncertainty had a weaker effect and was unable to account for the magnitude of the effect. The visual system is intrinsically uncertain about a target’s position (Hess & Hayes, 1994; Manjeshwar & Wilson, 2001; Semizer & Michel, 2017). Interestingly, we found that intrinsic position uncertainty better explained the results when the position uncertainty scales with the target size. This could happen if the level of intrinsic position uncertainty is constant in cortical space. Neuronal populations with larger receptive fields may tile visual space with the same overlap as small receptive fields; thus, their centers are further apart in visual space. Because of this, a fixed level of uncertainty in cortical space would give rise to variable levels of uncertainty in visual space depending on the size of the receptive fields encoding the stimuli. The same reasoning (the uncertainty being fixed in cortical space) can be applied when participants must detect a target presented in the periphery since receptive field sizes increase with eccentricity. Consistent with this idea, intrinsic position uncertainty has been previously shown to increase with eccentricity (Michel & Geisler, 2011). In the future, measuring the intrinsic uncertainty directly for different-size targets would be useful for testing whether scaling prediction holds.

Although intrinsic uncertainty can cause human efficiency to be higher under high levels of extrinsic uncertainty, limited attentional resources (another internal factor) may have the opposite effect. As the magnitude and number of extrinsic dimensions of uncertainty increases it may become more difficult to efficiently integrate the information across the dimensions, especially if the dimensions of uncertainty do not normally co-occur under natural conditions (like Hadamard set, see Burgess, 1985). Scale, orientation and position variability tend to co-occur under natural conditions, which helps to explain why we observed higher human efficiency relative to ideal when the level of extrinsic uncertainty was higher.

## General Discussion

Since the beginning of signal detection theory, the sum and max rules have been utilized to determine or approximate the ideal observer under various conditions (Peterson et al., 1954; Nolte & Jaarsma,1967; Graham et al., 1987; Ma et al., 2011). Here, we derived the optimal decision rule that combines the max and sum rules by inverting the generative environment. This generative environment consists of hierarchical set of exhaustive and mutually exclusive probabilistic events. This generative environment can represent many object identification tasks, such as object detection tasks where each category (like birds) involves many subtypes (like crows), covert visual search tasks where a target could appear in different orientations and scales, and face recognition tasks where any face can appear with many different expressions.

The conceptual framework presented here has three major restrictions. First, if many different targets exist in a single trial, the number of object categories increases combinatorically, as does computational demand. However, some of the demand can be alleviated by using previously computed likelihood ratios of subcategories through prefix sums. Secondly, the framework critically depends on the likelihood ratios of mutually exclusive events. However, sometimes, it is not possible to calculate the likelihood ratios exactly, and they are approximated by combining various cues. In that case, the equations will not provide the optimal performance, and one must be careful about how the approximation interacts with the decision-rule equations. Thirdly, even though we have provided the general equation when the context variables are unknown, the equation simplifies immensely if the context variables are known. A large background is usually available to the observer, often allowing the context variables to be estimated with good precision. However, in some cases, like visual search, there are no background regions where the target could not appear. Nonetheless, even under these circumstances, for natural scenes, basic background properties, like contrast and luminance, can be estimated from the target regions (Sebastian et al., 2017).

We have developed a method for effectively simulating the predictions of ideal and max observers. With a reasonably powerful CPU, we showed that the precomputations and later evaluation of psychometric functions for four dimensions of uncertainty took less than ten minutes. Thus, it is possible to handle representative simulations with even more dimensions of uncertainty with denser sampling. Since the precomputation is only done once, one can tolerate much longer precomputation times. With the large set of precomputed variables, it is possible to evaluate the performance for many tasks, levels of uncertainty, priors, and cost functions with different amplitudes and background contrasts.

During the development process, we also explored a sampling approach. Identifying the probability distribution of template responses for the basis condition is sometimes possible. In this case, the basis template responses might be generated through Monte Carlo simulations. Similarly, the parameters of corresponding probability distributions can be derived for various image properties using an expression similar to the one shown in the GitHub link. However, as the number of different conditions grows, it is very likely that each set of image properties will only correspond to a single trial. Thus, manipulating a large set of parameters to sample a single trial is inefficient. Hence, our method is more effective, especially for the high levels of uncertainty. On the other hand, it is possible to improve our simulation method. The computation times will improve with parallel processing and the use of a graphic processing unit (GPU), especially considering highly optimized libraries becoming available due to machine learning applications. However, it is best always to look for analytical simplifications to calculate model observers’ performance (e.g., Oluk & Geisler, 2023).

In this study, we also focused on comparing ideal observers with heuristic observers and differentiating these model observers by comparing them to human performance. First, we evaluated the heuristic max observers. Our results suggest that both template-energy and background-contrast normalization are important for approximating optimal performance and for explaining human performance (see Appendix, Figure 6b and OlukC Geisler, 2023). These normalizations specifically address the differences in the detectability across trials due to target scale and background contrast. Moreover, Ma et al. (2015) showed that not accounting for reliability differences while using the max rule yields some performance loss compared to the optimal performance and fails to account for human performance in general. Therefore, normalization (gain control) allows with max rule to achieve near-optimal performance. We suspect this benefit is not limited to the max operation, and thus normalization may be valuable for implementing heuristic strategies in general.

Here, we also showed that the SNN-MAX observer closely approximates the more sophisticated NN-MAX observer. Our finding is consistent with the recent findings of Zhang and Geisler (2024), where incorporating heuristic detectability maps into the decision rule is discussed more thoroughly. Thus, proper normalizations also make incorporating heuristic detectability maps into the decision process possible without loss of performance. Lastly, our previous work showed that under high levels of target amplitude and background contrast uncertainty, decision-making may only rely on a single criterion and avoid holding multiple criteria when responses are normalized by the background contrast (Oluk & Geisler, 2023). Taking all these together, we hypothesize that normalization’s important value is its capacity to make simple heuristic strategies work, which might help explain why normalization is so prevalent in the brain (Carandini & Heeger, 2012).

Given the simplification in decision processing provided by the normalization-dependent heuristic max models, it would be useful to see whether they can better explain human performance compared to ideal observers. Ma et al. (2015) showed that the model that addresses all the reliabilities differences, like the NN-MAX (in their formulation *max*_*d*_), performs slightly below optimal and is a slightly worse model of human behavioral data. Our behavioral data is consistent with SNN-MAX (equivalent to NN-MAX in our case) and the ideal observer, so our data cannot differentiate between these models. However, our findings provide insight into differentiating between these models. When we look at hit rate as a function of the target scale, the SNN-MAX observer’s predictions differ from the optimal performance because of the sharper decline (Figure 6b). Therefore, these models can make different predictions if we look at the pattern of hit and correct rejection rates even when their overall performance is similar (Figure 3). These differences are due to the distribution of subcategory likelihood ratios varying substantially across trials in which different targets are presented. For example, when the target is large, all subcategory likelihood ratios are highly correlated, whereas when the target is small, there are hardly any correlations between templates. These differences can occur within a single trial if the subcategories involve another layer of subcategories with different distributions of likelihood ratios. Therefore, future work can focus on designing tasks where the ideal and heuristic max observers make largely different predictions for hit and correct rejection rates to test which models best explain human data.

We also evaluated heuristic convolutional neural networks. We found that trained ResNet18 did not make any systematic errors compared to the optimal performance. However, training ResNet18 from scratch for a detection task under high levels of uncertainty requires a large amount of data (2.5 - 5 mil images), which could potentially be improved. Our findings suggest that reaching optimal performance in large uncertainty tasks is not trivial for CNNs. Thus, optimal performance can also provide an informative benchmark for deep learning models. For example, the definition of invariance is often not well-defined since the optimal detectability of object categories (or subcategories) is unknown, so human performance is commonly used as a goal for invariant detection (see DiCarlo & Cox, 2007; Pinto et al., 2008; Bart & Hegdé, 2012; DiCarlo et al., 2012; Han et al., 2020).

Here, we quantified how the optimal performance changes as a function of the target scale and showed that ResNet18 follows the same pattern, suggesting its detection is rotation invariant (no extra loss of sensitivity for any scale). Thus, future studies could evaluate the optimal confusion matrix for white noise backgrounds, especially for object detection tasks with many object categories (and subcategories) like CIFAR100. These optimal confusion matrices for object categories (and subcategories) will provide a rich pattern of results that can yield more insights into how CNNs function. Moreover, human performance can be measured to differentiate between predictions of various CNNs and the ideal observer to discover the underlying visual mechanisms.

Overall, we believe the conceptual framework and methods we developed in this study can help us understand the computational principles of many target identification tasks by providing a proper benchmark for heuristics observers. The ability to evaluate these image-computable models and optimal performance will allow us to make detailed predictions about the rich pattern of hit and correct rejection rates. These detailed predictions should make it possible to distinguish between ideal and heuristic observers (max, CNNs), providing important insight into the visual mechanism of many identification tasks.

## Appendix

### Derivation of the optimal decision rule

The maximum a posterior (MAP) estimate is given by the following:

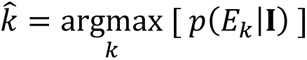

where **I** is the image that is presented on given trial. The posterior probability can be expressed as a sum over the context levels:

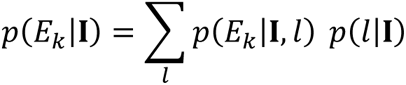

where *l* is an index to the set of vectors representing the relevant context variables. Note that we can also express *p*(*E*_*k*_|**I**, *l*) as *p*(*E*_*k*|*l*_|**I**). Next, we apply Bayes rule to the posterior probability conditionalized on the context:

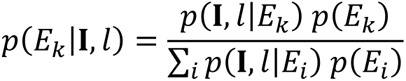

With some rearrangement, the posterior probability of object category k can be expressed as:

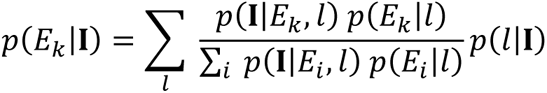

We divide both nominator and denominator by the likelihood of the target-absent object category (*p*(**I**|*E*_0_, *l*)) and define the likelihood ratio as:

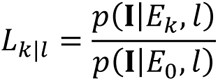

Now we can express the posterior probability of object category as:

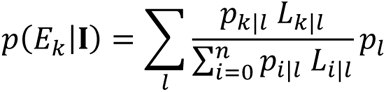

where *p*_*k*|*l*_ = *p*(*E*_*k*_|*l*) and *p*_*l*_ = *p*(*l*|**I**). Next, we express the likelihood ratio of an object category in terms of the sum of the likelihood ratios of its corresponding subcategories.

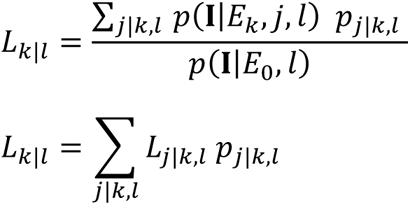

where *p*_*j*|*k*,*l*_ is the prior probability of subcategory *j* given the category and context. Finally, we can express the MAP estimate using the likelihood ratios:

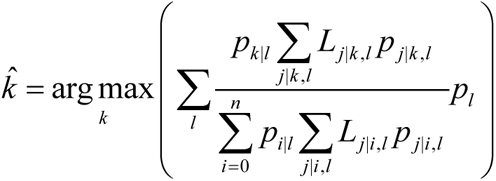

When the context can be estimated from the image (*p*_*ĵ*_ ≅ 1), we can ignore the sum over the context levels. Without the sum over context levels, the denominator does not change with the object category and thus has no effect on the maximum. Thus, the simplified equation is equal to the numerator. Finally, we also take the natural logarithm and rearrange the equation. The final simplified equation can be expressed as:

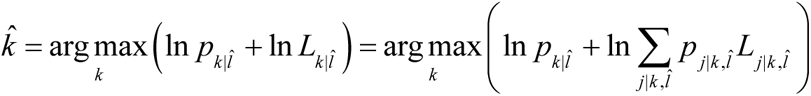

### Derivation of the optimal decision for additive targets in white noise

We start with Equation 2. The aim is to derive the likelihood ratio given the subcategory, category, and estimated context (*L*_*j*|*k*,_^*l*^^). For the additive target in white noise, the additive target is given as *a*_*j*|*k*_ **t**_*j*|*k*_ which is the product of an amplitude term (*a*_*j*|*k*_) and a target profile whose Euclidean norm is one, ǁ**t**_*j*|*k*_ǁ = 1. We assume the standard deviation of white noise is the known or precisely estimated (*σ̂*). Thus, we can express the likelihood ratio as multiplication of independent Gaussian variables:

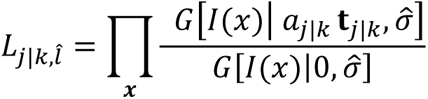

Writing the Gaussian explicitly and simplification of the same terms leads to:

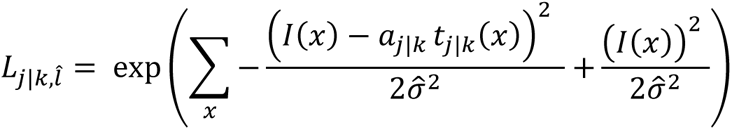

We expand the squared term and rearrange the equation:

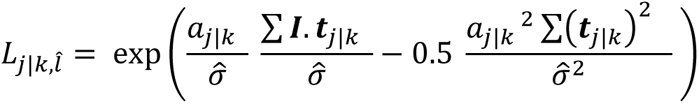

First, by definition ∑(***t****_j_*_|*k*_)^2^. Next, we define 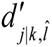 as *a_j_*_|*k*_/*σ̂*. We also define 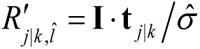. Finally, we replace the likelihood ratio in the general equation (Equation 2) with the derivation:

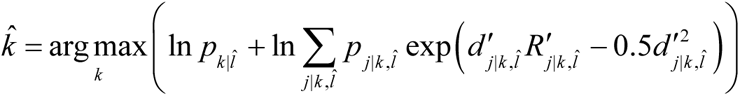

### Derivation of the heuristic max rule

First, we replace the sum operation with a max operation.

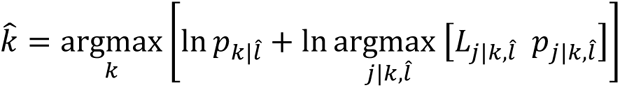

The natural logarithm is a monotonic operation that does not change the maximum, so we can change the order of the operations. Similarly, the logarithm of the prior probability of object category k given context does not depend on the subcategory; thus, it cannot change the maximum. There, we can change the order operation one more time.

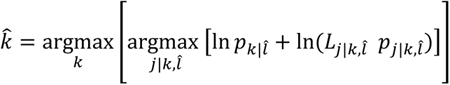

Next, we collect the logarithms together and merge the prior probabilities. Then, rearrangement of the equation leads to the Equation 4:

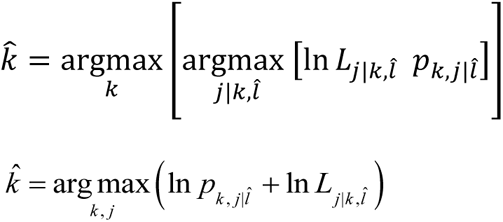

**Figure A1.**
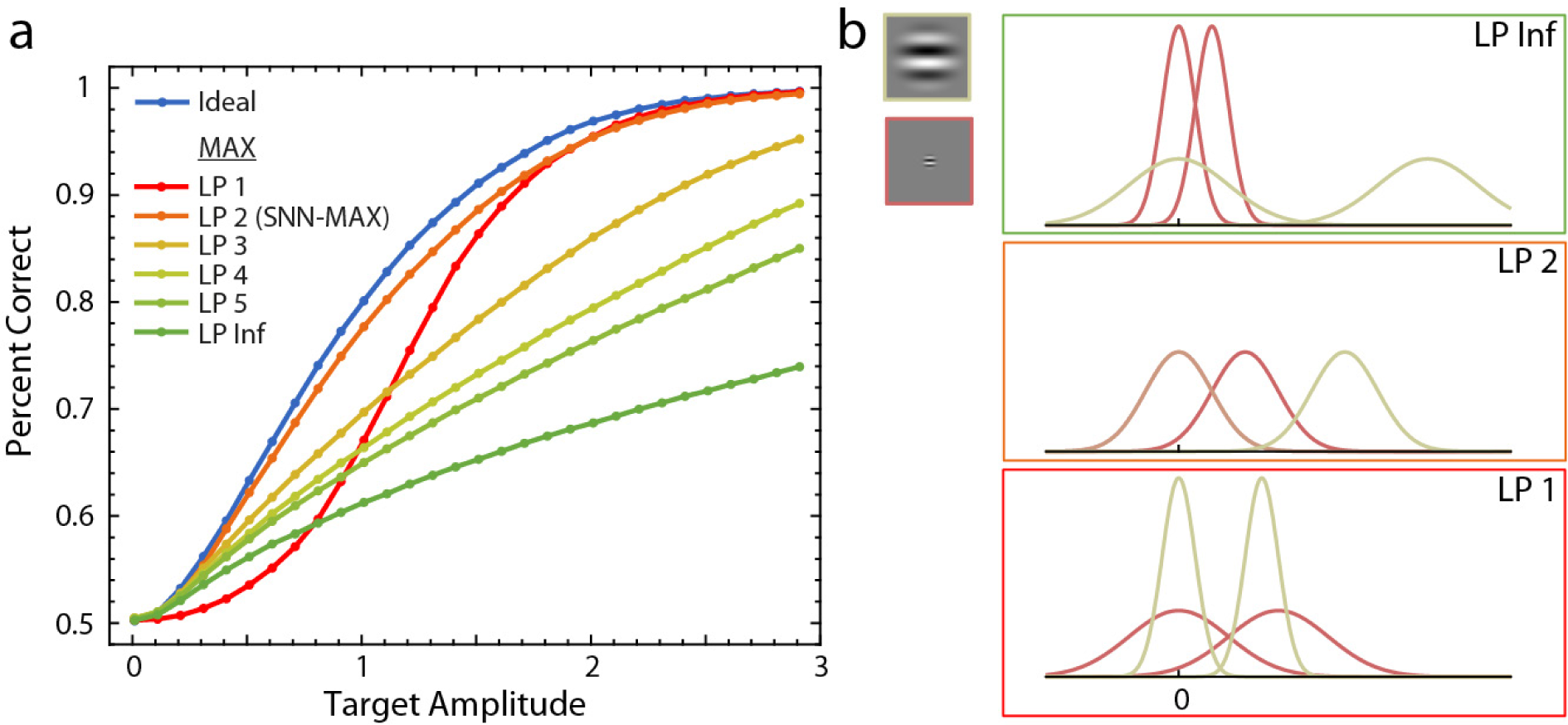
The comparison of the ideal observer with the family of MAX observers. **a** For MAX observers, template responses are divided with the *L*^*p*^ norm of the template for various levels of p. The criterion of the MAX observers is selected to maximize overall percentage correct, and overall percentage correct is shown as a function of target amplitude (A). **b** The illustration of the effect of normalization on the standard deviation of template response distributions for various levels of p. The template responses’ distribution is shown for two templates: a highly discriminable large target in green and hardly discriminable small target in light red. When the target is absent, distributions sit at zero. If the target that is the same as the template is presented, the mean is non-zero. The first panel shows the template responses when they are not normalized, the second panel shows them when they are normalized by the square root of the target’s energy of the target, and the last panel shows them when the normalization norm is one (*L*^1^).

**Figure A2.**
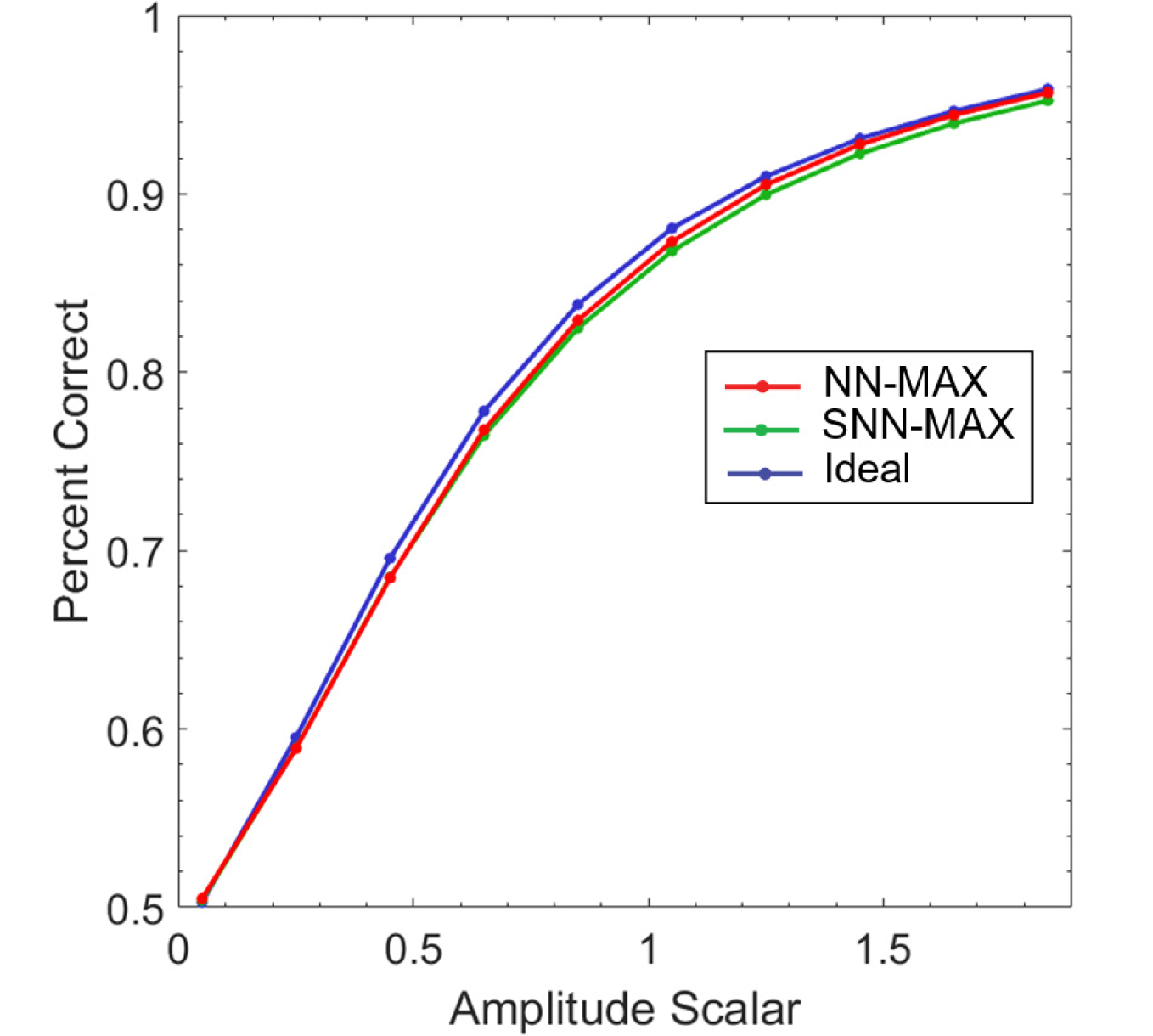
The comparison of the ideal observer with generalized normalized MAX observer and likelihood MAX observer. The NN-MAX observer picks the maximum likelihood ratio to make a decision and thus incorporates the relative d-prime map of various conditions. However, SNN-MAX observer picks the maximum normalized template response and thus assumes a flat d-prime map across conditions. The simulation parameters are exactly the same as the simulations shown in Figure 3. Even though NN-MAX performs slightly better than SNN-MAX, incorporating the d-prime map does not provide a large accuracy advantage.

**Figure A3.**
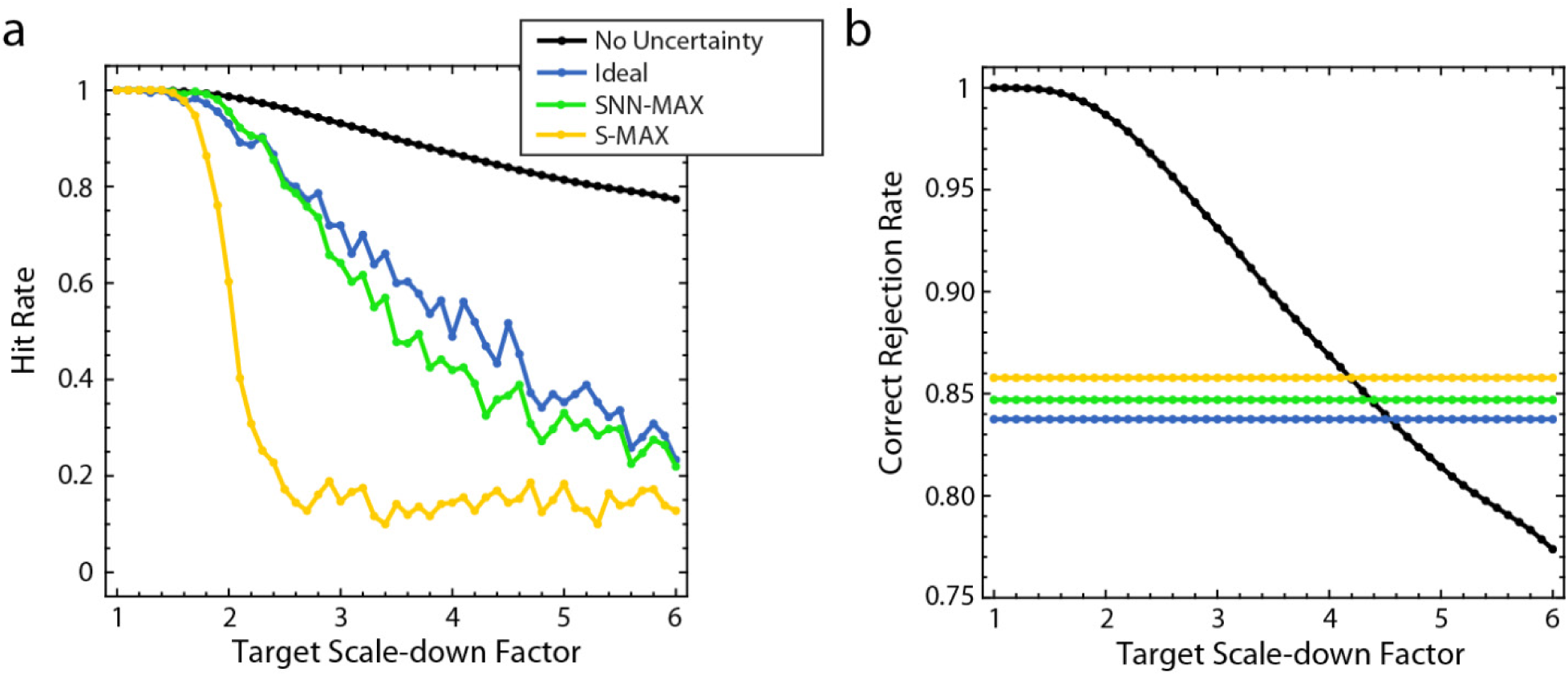
Hit and correct rejection rates of model observers under target orientation and scale uncertainty. **a** Hit rates as a function of target scale for three models under simultaneous scale and orientation uncertainty. The same rates are shown when there is no uncertainty about the scale and orientation. The target amplitude is picked such that the overall percentage correct is around 75 percent (threshold level). The expected hit rate patterns for different orientations are the same, so hit rates are averaged over orientation levels. Each point is calculated from 360 target present trials. When there is no uncertainty, all models’ predictions are the same. **b** Correct rejection rates as a function of target scale for three observer models under scale and orientation uncertainty. The same rates are shown when there is no uncertainty. When there is a combined target scale and orientation uncertainty, target absent trials do not differ systematically, so there is only a single correct rejection rate for a single uncertainty experiment.

**Figure A4.**
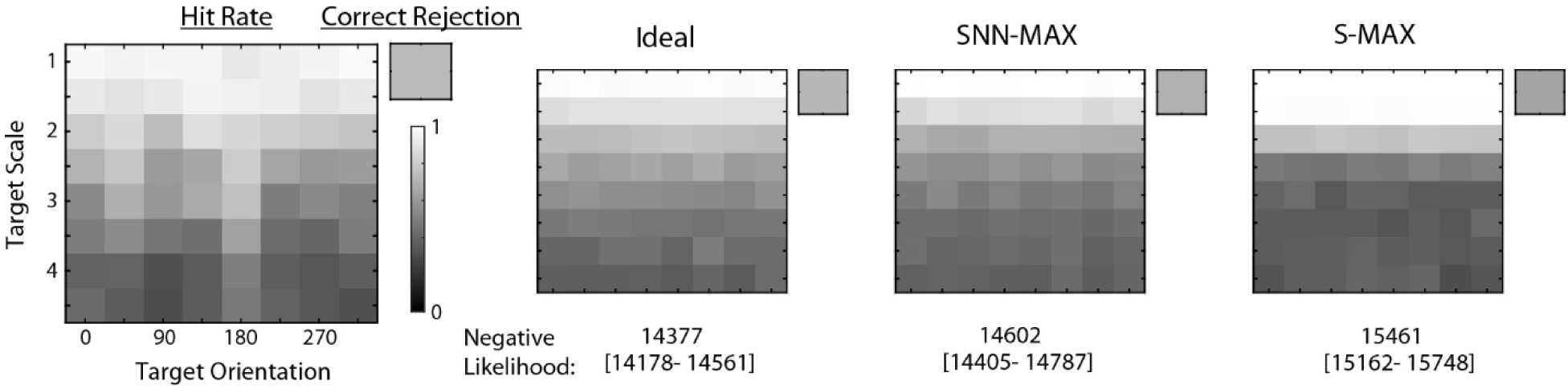
Data and model fits for experiment with high extrinsic uncertainty. The data matrix consists of an eight-by-eight hit rate matrix and a single correct rejection rate of the average participant. The average participant data is generated by aggregating the data of every participant. Grayscale is used to represent rates between 1 (white) and 0 (black). Fitted observer models’ hit and correct rejection rates are presented in the same structure, together with the negative log-likelihood associated with the fit and the 95 percent confidence intervals around it.

**Figure A5.**
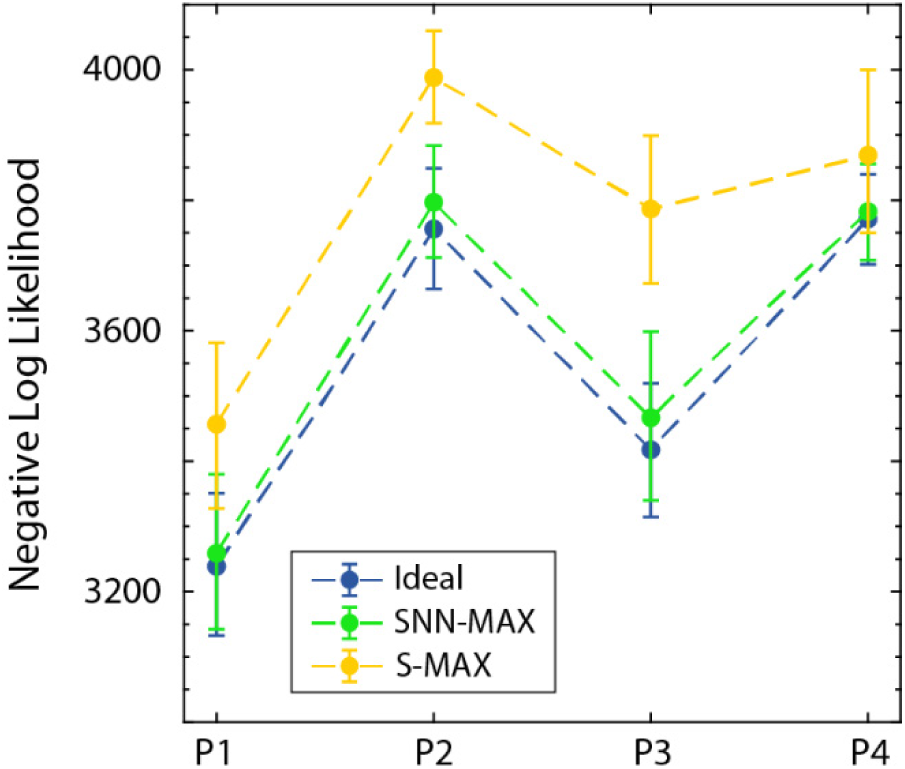
The goodness of fit measure (negative log-likelihood) for individual participants. Error bars show the 95 percent confidence intervals that are computed by fitting the bootstrapped data.

**Figure A6.**
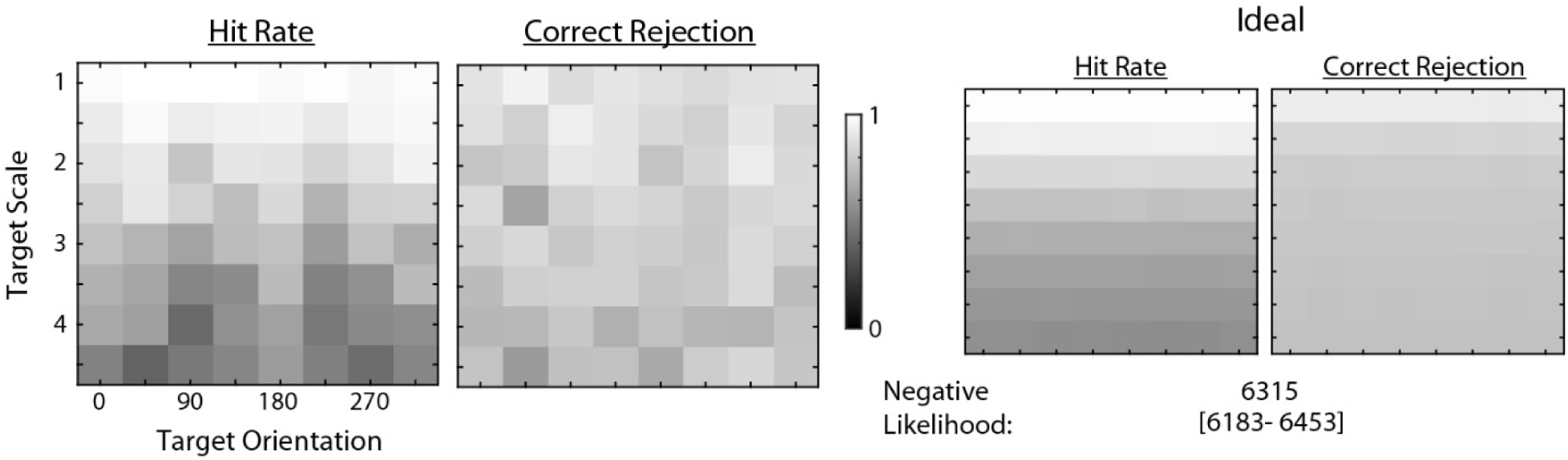
Data and model fit for experiment with low extrinsic uncertainty. The data matrix consists of an eight-by-eight hit rate matrix and an eight-by-eight correct rejection rate matrix of the average participant. Grayscale is used to represent rates between 1 (white) and 0 (black). The fitted model observer’s hit and correct rejection rates are presented in the same structure, together with the negative log-likelihood associated with the fit and the 95 percent confidence intervals around it.

